# Testing for changes in population trends from ecological count data

**DOI:** 10.1101/2025.01.08.631844

**Authors:** Liam Singer, Madleina Caduff, Andreas Füglistaler, Thierry Aebischer, Primaël Tabiti, Aimée Freiberg, Jens Ingensand, Christoph Leuenberger, Daniel Wegmann

## Abstract

1. Accurate, up-to-date knowledge on population trends are essential for effective biodiversity conservation and for evaluating interventions. Yet, estimating trends is often limited by sparse or noisy data, and many methods require long-term monitoring, delaying timely conservation actions.
2. We present birp, an R package that provides an accessible and fast Bayesian estimation frame-work designed to maximize power for detecting population trends. birp builds on classic count models, but extends them in six novel ways: i) it allows to explicitly test for changes in population trends under before-after, control-intervention, before-after-control-intervention or similar setups of arbitrary complexity, ii) it offers methods to identify and account for potential stochasticity and spatial variation in trends, iii) it allows to test and account for potential step changes, iv) it allows to test and account for overdispersion in the data, v) it implements appropriate priors, vi) relaxes the assumption of time-invariant detection probabilities through covariates and vii) models data from multiple survey methods jointly.
3. We illustrate the power of our tool through extensive simulations and show that changes in trends are accurately estimated under various designs, even with noisy or sparse data. We further demonstrate the usability of birp through two case studies: one illustrating the joint inference of trends from heterogeneous survey methods and irregular temporal sampling, and another showcasing its robustness under temporal subsampling. From these, we derive general guidelines for survey designs that maximize the power to detect trends.

## Introduction

Assessing population trends is an important goal of ecological monitoring, as they can serve as an early warning tool when threats emerge or intensify, such as increased poaching or the spread of a new disease. They also provide a direct way to assess the impact of interventions or policy changes by testing whether negative trends have slowed, halted, or reversed (Butchart et al., 2010; Wauchope et al., 2021; Volery et al., 2023). In both cases it is important to identify such changes in trends quickly, as biodiversity once lost can not be restored.

The counterfactual, which represents what would have happened in the absence of the intervention or the previous pressure regime, provides a conceptual standard for evaluating such changes (Khandker et al., 2009; Volery et al., 2023). Since natural conditions can rarely be truly replicated, experiments are often impractical in ecological contexts (Ferraro, 2009) and the counterfactual is commonly infered using quasi-experimental approaches (e.g. Geldmann et al., 2013; Terraube et al., 2020; Pellissier et al., 2020; Barnes et al., 2023; Langhammer et al., 2024). These include *control-intervention* (CI) designs, in which locations with and without the intervention are compared, *before-after* (BA) designs, in which trends before and after the intervention are compared at the same location, and *before-after-control-intervention* (BACI) designs that combine both approaches and are generally the most powerful (Wauchope et al., 2022)

A common approach to infer population trends from quasi-experimental data is to fit (non-)linear models to time-series counts of direct or indirect observations, accounting for stochasticity, variation in survey effort, and local conditions (Collen et al., 2009; Farmer et al., 2007; Wauchope et al., 2019). From such models, important change points, i.e. times at which the trend changes, can then be inferred *post-hoc* (Fewster et al., 2000). However, observational data results from stochastic processes modulated by local conditions, survey methods and survey efforts. To draw meaningful inference, they must thus either be perfectly standardized across surveys, or accounted for in the statistical analysis. Such time-series models are typically implemented by assuming exponential population change and fitting linear models under Poisson or negative binomial distributions (e.g. Williams et al., 2016; Moore and Barlow, 2011; Weir et al., 2009; Wauchope et al., 2019, 2022; Kéry et al., 2009). For example, in their work assessing the impact of protected areas on water birds, Wauchope et al. (2022) modeled site and year specific counts to estimate baseline population trends and any changes following the intervention. The intervention was considered impactful if including the post-intervention rate, that is, the change in trend after protection, significantly improved model fit.

We here develop Bayesian alternatives to such models, which are easy to use, applicable to a wide range of systems and arbitrary experimental designs, and are statistically powerful also in case of limited and noisy ecological data. In the following sections, we introduce the modeling framework, detail the birp software package, and present illustrative case studies.

## The birp package

birp provides a fast and accessible implementation of the Bayesian framework of Link and Sauer (1997a), and extends it to enable robust trend inference and change detection under heterogeneous sampling conditions. It handles multiple survey methods, accounts for variation in detection probabilities and survey effort, and accommodates arbitrary experimental designs, including BA, CI, and BACI.

For speed, birp was fully implemented in C++ and makes use of the dedicated library stattools. It is available through a Bitbucket repository at (link pending acceptance) as a stand-alone command line tool. For ease of use, it is also available as an R package birp on CRAN. A comprehensive vignette with detailed guidance and example applications complements the explanations provided below.

### Trend models

Link and Sauer (1997a) first realized that detection probabilities may not need to be known nor estimated if data can be stratified such that detection probabilities are time-invariant within each stratum. In their work, they used bird observers as examples: Consider two observers with very different detection abilities that both survey the same location. They may end up reporting vastly different observations. If the surveyed species doubled in numbers, however, both are expected to detect twice as many birds. As Link and Sauer (1997a) concluded, the total number of counts within a stratum is therefore reflective of the rate of detection, while their distribution across time is reflective of population trends. They further showed how this distribution can be modeled via conditioning under either a Poisson or negativ-binomial observation model.

birp implements and extends the framework of Link and Sauer (1997a) to estimate population trends and to formally test for changes in population trends as well as step changes from repeated count data collected across multiple locations potentially using multiple survey methods.

It partitions time into user-defined epochs. Within epoch *m*, trends in relative population abundance are modeled as exponential changes with rates *γ_m_*. At the turn between any two epochs *m* and *m* + 1, structural changes may occur: either the growth rate of the current trend may change, i.e. *γ_m_ ≠ γ_m_*_+1_, a step-change Δ*_m_* (i.e. a jump in relative abundance) may occur, or both. The user can arbitrarily group locations that may differ in either parts of their entire trends. In the case of a classic BACI design, for instance, two groups of locations would exist, those with and those without the intervention, with both sharing the trends in the first epoch prior to the intervention, but not after that. birp allows for arbitrary designs consisting of arbitrarily many groups and epochs; see Figure 1 for illustrations. For each design, birp the provides Bayesian estimates of the exponential rates and the strengths of step changes. See the Supplementary Information for a detailed description of the full model.

**Figure 1:**
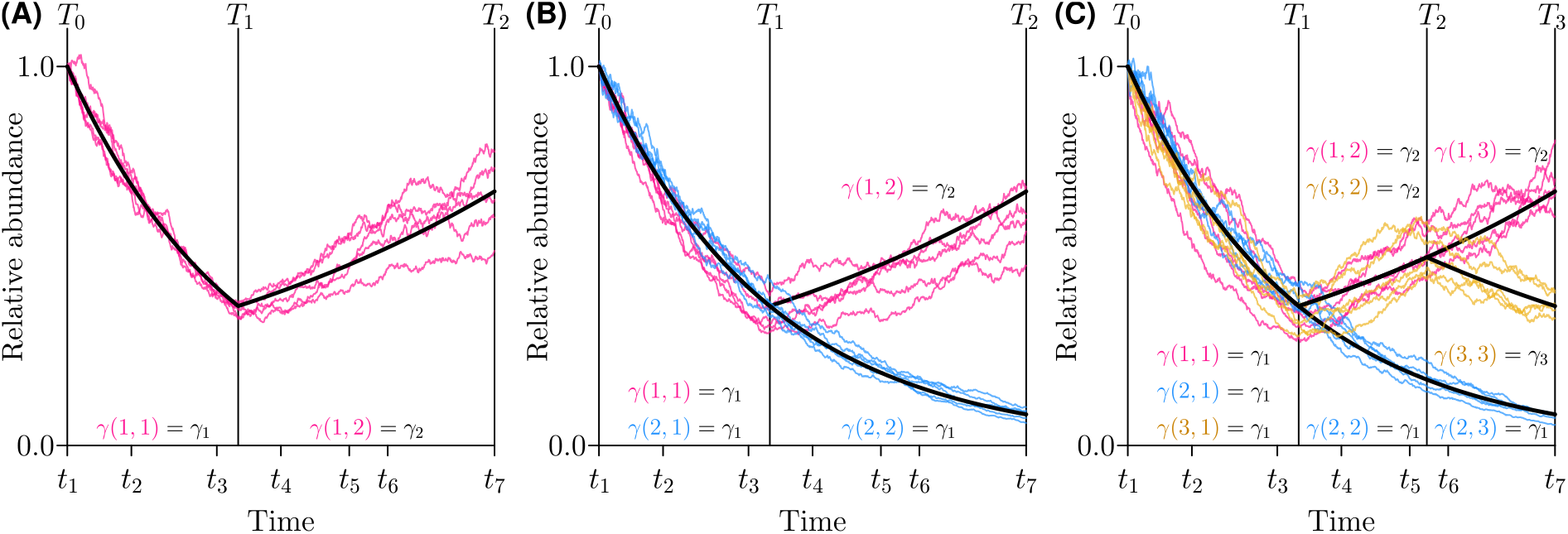
Illustrations of birp models with seven survey times under different designs. Black vertical lines mark epoch boundaries. The deterministic population trend is shown in black and possible stochastic realizations in color. We used three different rates *γ*_1_ = *−*0.25, *γ*_2_ = 0.1 and *γ*_3_ = *−*0.1. The relevant exponential rate for locations of group *g* in epoch *m* is denoted by *γ*(*g, m*). (A) A canonical before-after design with two epochs. A canonical before-after-control-intervention (BACI) design with two epochs and two groups: a control group (blue) and a group (pink) affected by an intervention at time *T*_1_. (B) A complex BACI design as in B but with an additional group (gold) affected by an additional intervention at *T*_2_.

Two main model variants are implemented: The deterministic model assumes that all locations within a group follow the same exponential trend in each epoch and experience the same step changes (black lines in Figure 1). In contrast, the stochastic model allows for some variation across the locations within a group modeled as geometric Brownian motion and infers the degree of variation (the volatility) from the data. For both variants, birp allows for both Poisson and negative-binomial observation models to allow for potential overdispersion in the data and implements the the non-informative Jeffreys prior initially proposed by Aebischer et al. (2020) and extended her for both trend and step-change parameters. Inference is performed in a Bayesian framework using MCMC sampling. See the Supplementary Information for a detailed description of the inference scheme.

Importantly, the framework does not require a strictly regular or complete temporal sampling design: survey occasions may be irregular and sampling gaps are naturally accommodated through the likelihood formulation. In addition, data collected under different sampling methods such as track counts, camera trap images, distance sampling or metabarcoding sequences can be analyzed jointly, thereby increasing precision by borrowing strength across data sources. Systematic differences between survey conditions that affect detectability such as weather, time of day, or season can also be incorporated as method-level strata, when relevant. The assumption of time-invariant detection probabilities within strata can further be relaxed by modeling detection and survey effort as functions of covariates, allowing detection to vary over time, albeit at the cost of reduced statistical power. We return to this assumption and their practical implications in the Discussion.

### Data input and analysis workflow

birp infers trends jointly from multiple survey methods. To prepare data for analysis, the data for each survey method should be provided either as a separate dataframe within R or as a CSV file listing location identifiers, survey times, observed counts, survey efforts (or effort-related covariates), and the identifier of the group to which this survey belongs. Once formatted, the data are converted into a birp_data object using the function birp_data_from_data_frame or birp_data_from_file. The trend estimation is done with the main function birp(). By default, birp() assumes a Poisson likelihood, but a negative-binomial model can be fitted if overdispersion is present by specifying negativeBinomial = TRUE. Analogously, users can specify stochastic models (stochastic = TRUE) or multiple epochs (timesOfChange). Estimates are then available as objects of type birp, which can be visualized using built-in plotting functions. A full step-by-step tutorial, including input data and explanations of all modeling options, is available in the package vignette.

### Model selection and analysis tools

The package includes specific tools to guide analysis and interpret results:

#### Distinguishing between Poisson and negative binomial models

We implemented a simulation-based approach to check whether the inferred overdispersion differs significantly from what is expected under a Poisson model, thus guiding the choice between Poisson and Negative Binomial models. This test is available directly to the user via the function assess_NB.

#### Trend Visualization

The package provides several functions for visualising estimated trends. The function plot generates density plots of the posterior distributions of the parameters *γ_m_* and Δ*_m_* regarding exponential rates and step-changes, visualizing the uncertainty in the estimated parameters. The function plot_trend shows the posterior trend of relative densities over time, including credible intervals and optional epoch or survey markers. Finally, plot_epoch_pairs allows comparison of posterior densities between pairs of epochs, summarising shifts in estimated abundance between specific time periods.

#### Hypothesis Testing

The estimated parameters are informative regarding highly relevant questions for ecological monitoring:

- Is a population increasing or decreasing during epoch *m*? Given the full data ***n***, this question is answered by the posterior probabilities ℙ(*γ_m_ >* 0|***n***) or ℙ(*γ_m_ <* 0|***n***) for the population with the function prob_trend.
- Did a population trend change at the turn of epochs *m* and *m*+1? This question has so far only been addressed indirectly and under parameter rich models (e.g. Fewster et al., 2000). It can readily be answered by determining the posterior probabilities ℙ(*γ_m_ > γ_m_*_+1_|***n***) or ℙ(*γ_m_ < γ_m_*_+1_|***n***), which are obtained with the function prob_trend_diff.
- Do the rates *γ* and *γ^′^* differ between two groups? This question, which arises in both CI and BACI designs, and is answered by the posterior probabilities ℙ(*γ > γ^′^|****n***) or ℙ(*γ < γ^′^|****n***). These are also obtained with function prob_trend_diff.
- Did a step change occur at the turn of epochs *m* and *m* + 1? This questions is answered by the posterior probabilities ℙ(Δ*_m_ >* 0|***n***) or ℙ(Δ*_m_ <* 0|***n***). These are obtained with function prob_step.

##### Performance against simulations

To evaluate the performance of birp, we conducted extensive simulations across a range of ecological monitoring scenarios, including under the canonical BA, CI and BACI designs. We systematically varied the number of survey time points, number of locations, observation effort, and levels of overdispersion or stochasticity. Trend-detection power was generally high and increased with more survey time points, more locations, stronger underlying trends, and lower overdispersion or volatility (Figure SS1 and SS2). We also explored cases such as local extinctions (all-zero counts) and delayed intervention effects, and found that birp retained robust performance in most cases, with reduced power primarily in scenarios involving long delays between intervention and effect (Figure SS3). Power to detect step changes was also generally high, and increased with more survey time points and larger step sizes (Figure SS4). However, we noted that step changes could not always be reliably distinguished from rate changes. Additionally, we introduce a simulation-based approach to test the adequacy of the Poisson assumption, enabling informed model selection that balances power and false-positive control in the presence of overdispersed data (Figure SS5). Detailed simulation setups and results are provided in the SI.

## Case Study 1: Temporal downsampling

We reanalyzed temporal subsets of 35 years of Tasmanian devil (*Sarcophilus harrisii*) transect counts from six bioregions of Tasmania (1985–2019) (Driessen and Hocking, 1992; Cunningham et al., 2021) to test for changes in population trends following the emergence of devil facial tumour disease (DFTD) (Hawkins et al., 2006; McCallum et al., 2007). For each bioregion, we designated the first survey in which DFTD appeared in at least half of the transects as year 1, and set the time of change at the midpoint between that survey and the previous one (*T*_1_ = 0.5). We downsampled the data to use five years before and one to eight years after the regional DFTC outbreak for every bioregion. We treated each transect as a separate location, ran the analysis using a single group, and selected the appropriate likelihood model (Poisson or negative binomial) for each bioregion using *assess_NB*, applying the Poisson model when *p >* 0.05.

Counts were highly variable across years and regions (Figure 2A). Using birp, we estimated separate rates before (*γ*_1_) and after (*γ*_2_) *T*_1_, and computed the posterior probability that the population trend became more negative (ℙ(*γ*_1_ *> γ*_2_); Figure 2C). Strong support for a post-DFTD decline was found in bioregion *Ben Lomond* (ℙ *>* 0.99 with three or more surveys after *T*_1_) and *Northern Slopes* (ℙ *>* 0.95 with 3–4 surveys). *Central Highlands* and *Southern Ranges* required more post-change surveys to detect a decline, reflecting delayed or gradual effects. *South East* showed an immediate drop after *T*_1_, but partial recovery in later surveys, while *King* showed little evidence of decline, likely because the disease had not yet fully spread to this bioregion (Lazenby et al., 2018). These results are both compatible with a visual inspection of the data as well as with previous analyses (Lazenby et al., 2018; Volery et al., 2023) and demonstrate that birp can detect population trends even from short time series. Further technical details and extended discussion are provided in the SI.

**Figure 2:**
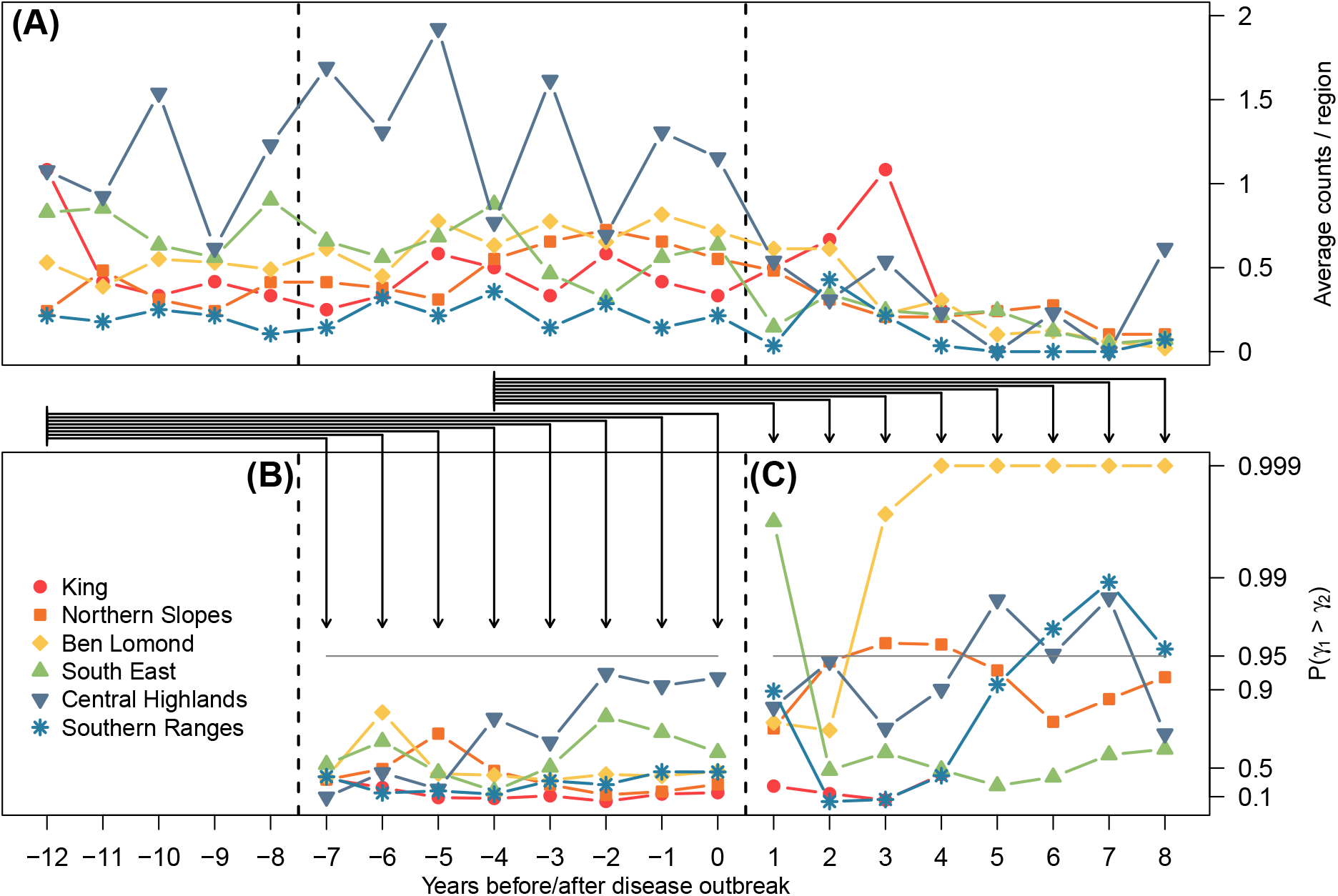
Analysis of Tasmanian devil transect count data. (A) Average counts per region over 21 years, aligned by the estimated time when the disease was present at half or more of the locations within each bioregion, denoted by year 1. (B,C) Posterior probabilities that the first-epoch rate parameter (*γ*_1_) exceeded the second-epoch rate (*γ*_2_), shown on a logarithmic scale for two potential times of change (dashed vertical lines): year 0.5 (C) and eight years earlier (B). Each analysis used five years before the time of change and one to eight years after, as indicated by the arrows. The horizontal lines mark the 95% certainty threshold. The x-axis applies to all panels.

As a control, we repeated the analysis using a period of eight years before the outbreak, when no known major ecological changes occurred (Figure 2B). No spurious changes were detected in any region, confirming that birp accurately distinguishes true population trends from random variation.

## Case Study 2: Heterogeneous survey methods and irregular sampling

We tested for changes in population trends following active law enforcement in the Aire de Conservation de Chinko (hereafter *Chinko*) in the eastern Central African Republic, a remote region with limited historical biodiversity data (Aebischer et al., 2017). While formally protected, active management only began in 2016. We reanalyzed track count and camera trapping data for four apex predators and six large prey species from Aebischer et al. (2020); Ait Kaci Azzou et al. (2021), supplemented with additional data collected under the same protocol. The dataset spans 2012–2024, with some years missing due to logistic constraints; birp explicitly handles such sparse sampling.

We fitted a deterministic model with a single group, two epochs (*γ*_1_, *γ*_2_), and a change point at 2017, chosen as the first year of full active law enforcement. We determined the approriate model (Poisson or NB) for every species using the function *assess_NB* and used the Poisson model when *p >* 0.05. Track count surveys after 2022 were treated as a separate method due to a change in the main observer. Additional technical specifications are listed in the SI.

For most species, our inference indicated a currently positive growth rate and a positive change in trend following law enforcement, with generally high posterior support (ℙ(*γ*_1_ *< γ*_2_ *| n*)), although uncertainty was greater for species with fewer observations (Figure 3). Our results are in good agreement with those of Aebischer et al. (2020), despite methodological differences: First, we analysed the data under the negative binomial model rather than the Poisson model for most species to account for the overdispersion present in the data. Second, we treated each road segment as an individual location and camera trap observations from dry and wet seasons as different methods, choices which better account for spatial and temporal variation in detectability. Third, we directly tested for a change in trends, rather than inferring trends individually for the different epochs. The somewhat lower current growth rates observed for some predators (especially lions and leopards) may be due to our use of much longer time series in the second epoch, over which trends may be more moderate than what was observed right after law enforcement was enacted. Overall, the analysis illustrates that birp handles irregular temporal sampling and mixed survey methods well, and provides reliable and robust inference even in data-limited systems. Detailed results for each species are available in the SI.

**Figure 3:**
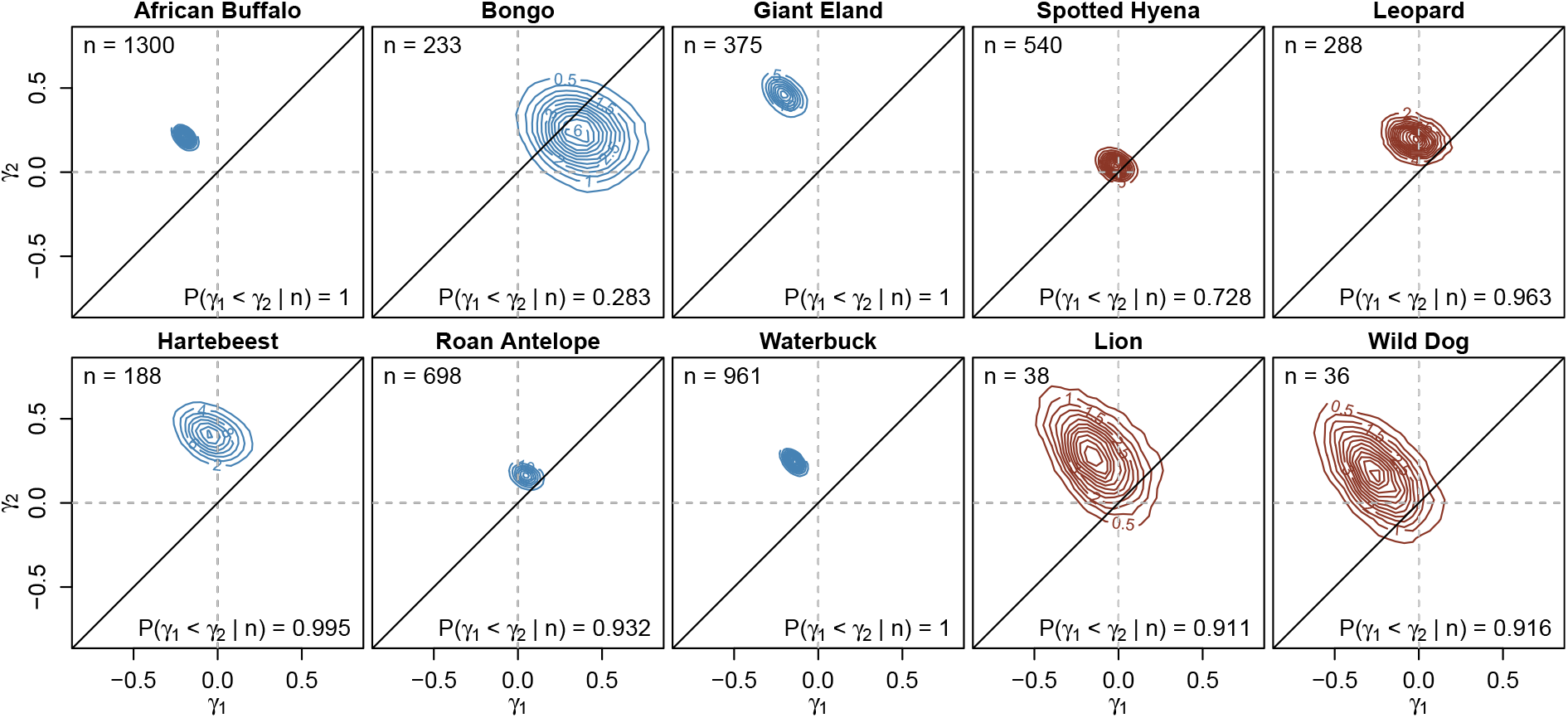
Analysis of survey data from the Chinko. Shown are two dimensional posterior densities for the trends before (*γ*_1_, x-axis) and since active law enforcement was enacted (*γ*_2_, y-axis) for four apex predator species (right, brown) and six of their potential prey species (left, blue). The total number of observations is shown in the top-left corner; the posterior support for a more positive trend since law enforcement was enacted in the lower-right corner.

## Discussion and Practical Considerations

### Population trends vs. population trajectories

The terms population trend and population trajectory are sometimes used synonymously, although their meaning differs with potential implications for conservation management. We here refer to Link and Sauer (1997b), who briefly discussed this terminology in their commentary and proposed to use the term *trajectory* to “describe the pattern of population fluctuation” and the term *trend* to “represent an average rate of change over a specific time interval”. Thus, a trend, according to this definition, is a smoothed trajectory (Link and Sauer, 1997b). Following this, it is easy to recognize that a population may show fluctuating trajectories while overall exhibiting a stable trend. Conversely, a declining population could temporarily show a positive trajectory as a result of random fluctuations. Hence, being able to distinguish longer-term trends from shorter-term variation in trajectories is important, yet these definitions are highly context and species dependent and there is no universally valid definition for when a trajectory turns into a trend. For these reasons, there is no consensus on the time span needed to reliably estimate population trends (e.g. Brashares and Sam, 2005; Maclean et al., 2013; Embling et al., 2015). As shown through simulations and real data applications, birp may reliably identify changes in abundance from much fewer data points. Whether those inferred changes reflect random fluctuations or longer-term trends will be up to the experimenter to judge. The risk of misinterpreting random fluctuations as trends is, of course, greatly reduced when surveys were conducted over longer times, but also if many locations were surveyed such that random fluctuations effectively cancel out. We argue that from a biodiversity conservation perspective, it may be crucial to act even if a sharp change in trends was inferred from short time series only.

### Time-invariant detection probabilities and survey design

The framework initially put forth by Link and Sauer (1997a) and implemented in birp assumes that detection probabilities are time-invariant within strata. While detection probabilities likely vary greatly between locations and methods (e.g. Thompson and La Sorte, 2008; Diefenbach et al., 2003), it is often possible to stratify observed data to meet this assumption. For track-count surveys, for instance, detection probabilities were reported to differ between soil types (e.g. Funston et al., 2010). Similarly, detection probabilities of camera traps are known to vary greatly with habitat structure (e.g. Ait Kaci Azzou et al., 2021; Hofmeester et al., 2019). However, such variation is readily accounted for if the same locations are repeatedly surveyed. In our analysis of track count data (Case Study 2), for instance, we stratified the data by segments of the surveyed road network. While there may also be variation in detection probabilities within a segment, that variation is of no concern if the segments are defined such that they are always surveyed in full. For the camera trapping data, we similarly only considered locations at which camera traps were mounted at multiple survey times, if possible at the very same tree. Detection probabilities likely also varies across methods, observers or even across devices. For trend inference, data is thus ideally collected such that the very same method, observer and even device is employed at a specific location across surveys, as illustrated in Figure 4.

**Figure 4:**
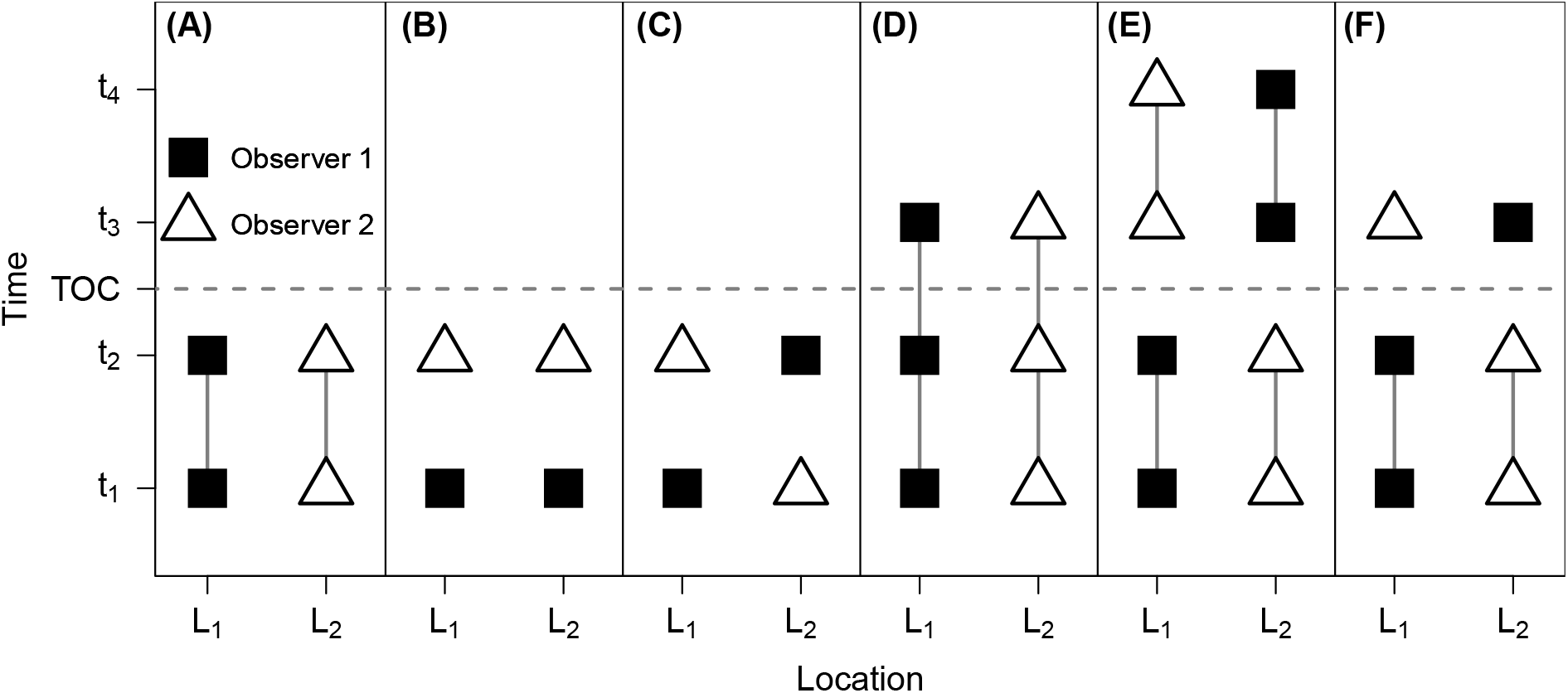
Different experimental designs that allow (panels A, D and E), partially allow (panel F) or do not allow (panels B and C) for stratification. Two different observers (or survey methods) are indicated with a black square and a white triangle, respectively. Lines connecting the observers across survey times indicate surveys that are comparable in terms of detection probabilities and thus form a stratum. The horizontal dashed line indicates a potential time of change (TOC).

Detection probabilities may further vary due to other factors such as the weather condition, the season, or the time of the day. Some of these factors can also be readily stratified. If surveys were conducted at night and day, for instance, they should be considered as different “methods”, even if conducted by the same observer, and analogously for different seasons or weather conditions. As the number of such factors grows, however, so will the resources required for planning and logistics to ensure enough comparisons are available within each stratum. It may thus be necessary to merge similar strata, such as, for instance, observers with similar skills. Alternatively, birp also allows to model variation in detection rates through covariates, although additional parameters then need to be estimated. Accordingly, users should decide based on the structure and quality of their data whether the simpler assumption of constant detection or a more complex covariate-based detection model is most appropriate. Importantly, the covariates are not required to explain absolute detection probabilities, they only need to account for relative differences in detectability.

We also note that in some reports, detection rates were found to vary with abundance, e.g. that movement, call rates, or space-use changes with local density (Warren et al., 2013; Neubauer et al., 2022; Broadley et al., 2019). This may lead to underestimation of actual trend changes. While correcting for such an effect would require explicit knowledge on the relationship between abundance and detection, we expect that birp remains robust for identifying the overall direction of population trends also in such case, though the estimated magnitude of change may be somewhat affected.

### The Poisson assumption

Ecological data is often overdispered in that it violates the Poisson assumption of mean equal variance. To maximize statistical power, we here implemented both Poisson and negative binomial models, as well as a simulation-based test to guide users towards the appropriate model for their data. However, we note that zero-inflation, a particular case of overdispersion that results from a sequence of non-observations due to the absence of a species at a specific location (Blasco-Moreno et al., 2019), does not affect the models presented here: If a species is truly absent from a location, no counts were obtained across all years surveyed. Such locations bear no information regarding trends and, to avoid unnecessary calculations, are omitted.

### Changes in trends vs. step changes

In our simulations, we observed that birp can not always distinguish between step-changes and changes in trends. If data from a single time point after the change was avilable, the two scenarios become truly indistinguishable, but the distinction appears to remain challenging for short time series in general. We therefore recommend that users should run the inference allowing for only one the two effects at the time and argue that the distinction between these two scenarios is unlikely relevant for short time series as both models are powerful to detect recent changes, regardless of the underlying cause. Using simulations, we for instance found that a model inferring changes in trends accurately identifies step-changes under a wider range of cases (see Supplementary Information). For longer time series, the distinction between step changes and changes in trends becomes more relevant, of course, and we recommend that users run models allowing for both to occur, and to then simply models if there is limited posterior support for one of the two processes.

### Conclusions

We here introduced birp, a Bayesian framework for inferring population trends and changes in trends from ecological count data. Our simulations and real-world applications demonstrate that the model performs robustly and remains effective even when data are sparse, noisy, or collected under irregular sampling designs. In the future, the method could readily be extended to also infer the timing of epoch boundaries or even the number of epochs, for which data if multiple species analyzed jointly might be particularly informative. While current implementation offers a single-species model only, we do note that data from multiple species can be analyzed under the assumption that they share the same trends.

## Acknowledgements

We would like to thank the African Parks Network and its hard-working staff. Special thanks go to the ecological research team in the Chinko: Modeste Bambalo, Narcisse Bangali, Alain Gbando, Emmanuel Lango, Fernand Magraba, Vivien Pandi and Junior Wakassa. This work was supported by a Swiss National Foundation grant 310030_208154 to DW.

## Author Contributions

LS, MC, DW and CL conceived the ideas and designed methodology; TA and PT collected the data; MC and LS implemented the methods, LS, MC, AF and JI analyzed the data; LS, MC and DW led the writing of the manuscript. All authors contributed critically to the drafts and gave final approval for publication.

*Inclusion statement* Our study brings together authors from two different countries, including scientists based in the country where data collection for this study was carried out. All authors were engaged early on with the research and study design to ensure that the diverse sets of perspectives they represent was considered from the onset. Whenever relevant, literature published by scientists from the region was cited; efforts were made to consider relevant work published in the local language.

## Conflicts of interest

The authors declare no conflict of interest.

## Notes

### Competing Interest Statement

The authors have declared no competing interest.

### Summary of Updates

This version reflect a major revision based on detailed reviewer comments. In particular, the manuscript is now formatted as an Application Note and considerably shorter with many details now found in the supplementary material.

