## Supplementary Material for "Testing for changes in population trends from ecological count data"

#### Modeling population trends

We here present a Bayesian methods to infer population trends from count data obtained with (potentially) multiple survey methods such as track counts, camera trap images, distance sampling or metabarcoding sequences. The methods build on classic Poisson and negative binomial models for ecological count data (in particular Link and Sauer, 1997; Kéry et al., 2009), for which we introduce both a deterministic and a stochastic variant that allow to directly test whether population trends changed at specific times, while integrating over the uncertainty associated with absolute densities and detection probabilities.

##### Deterministic approach

We assume that the data were obtained from $J$ locations, which can be grouped into $G$ groups such that all locations within group $g$ share the same trend. Following others (e.g. Williams et al., 2016; Moore and Barlow, 2011; Weir et al., 2009; Kéry et al., 2009; Wauchope et al., 2022), we model these trends in population sizes exponentially with rates $\gamma_l$, where a positive and negative value indicates an increase or decrease in abundance, respectively. Similar to e.g. Wauchope et al. (2022), we allow for structural changes to happen at fixed times $T_m, m = 1, \ldots M-1$ at which the rate parameters may change, a step-change may occur, or both. The trends are thus divided into $M$ epochs, of which epoch $m$ runs from $T_{m-1}$ to $T_m$.

Let $\boldsymbol{\gamma} = (\gamma_1, \ldots, \gamma_L)$ denote the vector of all rates, which may be shared between groups, and let $\gamma(g, m) \in \boldsymbol{\gamma}$ denote the relevant rate for locations of group $g$ during epoch $m$. Similarly, let $\boldsymbol{\Delta} = (\Delta_1, \ldots, \Delta_D)$ denote the vector of all step changes, also potentially shared between groups, and let $\Delta(g, m)$ denote the step change relevant for locations of group $g$ at time $T_m$, i.e. between epochs $m$ and $m + 1$. Figure **??** shows visualizations of this model for several cases, including a canonical BACI design with two epochs and two groups, of which the intervention group has two potentially different rates $\gamma(1, 1)$ and $\gamma(1, 2)$ for the two epochs, while the control group has the former rate in both epochs ($\gamma(2, 1) = \gamma(2, 2) = \gamma(1, 1)$, Figure **??**B). However, the approach described here allows for an arbitrary number and combination of groups, epochs and rates (e.g. Figure **??**C).

Our method then infers the full vector of parameters $\boldsymbol{\theta} = (\boldsymbol{\gamma}, \boldsymbol{\Delta})$ consiting of the rates $\boldsymbol{\gamma}$ and step changes $\boldsymbol{\Delta}$ from observations obtained during multiple surveys $k = 1, \ldots, K$ conducted at known times $t_k$. The data does not contain information about changes in the rates before the first ($t_1$) or after the last ($t_K$) survey. We therefore assume $t_1 < T_m < t_K$ for $m = 1, \ldots, M - 1$ and define $T_0 = t_1$ and $T_M = t_K$ (Figure **??**). Note also that in case a survey was conducted exactly at the turn of an epoch (e.g. $t_3 = T_1$), we assume that a potential step change at that time occured prior to the survey. We do therefore not consider step changes at $T_0$.

Let us denote by $\varphi_j(t_k, \boldsymbol{\theta}) = N_{j,t_k}/N_{j,t_1}$ the abundance at $t_k$ relative to that at the first time point $t_1 = T_0$ for location $j = 1, \ldots, J$. For example, $\varphi_j(t_k, \boldsymbol{\theta}) = 1$ implies the same abundance as at time $t_1$ and $\varphi_j(t_k, \boldsymbol{\theta}) = 0.1$ a reduction by 90%. Under the deterministic exponential model proposed above, $\varphi_j(t_k, \boldsymbol{\theta}) = e^{\gamma(g,1)t_k}$ for any location $j$ in group $g$ and for any $t_k$ in the first epoch $T_0 \leq t_k \leq T_1$. At $T_1$, step change $\Delta(g, 1)$ may occur and the rate $\gamma(g, 2)$ will take over. For a survey time $t_k$ in the second epoch $T_1 < t_k \leq T_2$, therefore, $\varphi_j(t_k, \boldsymbol{\theta}) = e^{\gamma(g,1)(T_1-T_0)+\gamma(g,2)(t_k-T_1)+\Delta(g,1)}$ for any location $j$ in group $g$.

More generally, and denoting by $g(j)$ the group to which location $j$ belongs,

$$\varphi_j(t_k, \boldsymbol{\theta}) = \exp\left[\sum_{m=1}^{M} \rho_m(t_k)\gamma(g(j), m) + \sum_{m=1}^{M} \mathbb{1}_{t_k \geq T_m} \Delta(g(j), m)\right], \tag{1}$$

where the indicator function $\mathbb{1}_{t_k \geq T_m}$ is one if survey time $t_k$ was after time $T_m$ and where we introduced the factors

$$\rho_m(t_k) := \begin{cases} T_m - T_{m-1} & \text{if} \quad T_m \leq t_k, \\ t_k - T_{m-1} & \text{if} \quad T_{m-1} < t_k < T_m, \\ 0 & \text{if} \quad t_k \leq T_{m-1}, \end{cases} \tag{2}$$

which indicate whether the sampling time $t_k$ lies after, within or before epoch $m$, in which case the rate $\gamma(g(j), m)$ was active for the full length of the epoch, the time passed since the start of the epoch, or not at all, respectively.

#### Stochastic approach

Stochastic models allow to account for differences in realized population trajectories between survey locations that are part of the same group. Many existing stochastic trend models use integer abundances in discrete time (e.g. Wang, 2007; Pedersen et al., 2011; Boyd et al., 2018; Sollmann et al., 2015; Dennis et al., 2006; Hostetler and Chandler, 2015; Buckland et al., 2004; De Valpine and Hastings, 2002), often with considerable realism such as density dependence or migration. To maximize the statistical power to infer trends from small data sets, we here follow Dennis and Ponciano (2014); De Valpine and Hastings (2002) and take a continuous approach.

A natural stochastic variant of the deterministic, exponential model described above is geometric Brownian motion (see Karlin and Taylor, 1981, Ch. 15), which describes the stochastic trend $\tilde{\varphi}_j(t, \boldsymbol{\theta})$ specific to location $j$ using the stochastic differential equation:

$$d\tilde{\varphi}_j(t, \boldsymbol{\theta}) = \gamma_{g(j)}(t)\tilde{\varphi}_j(t, \boldsymbol{\theta})dt + \sigma\tilde{\varphi}_j(t, \boldsymbol{\theta})dW(t),$$

where the volatility $\sigma$ is the only additional parameter,

$$\gamma_{g(j)}(t) = \sum_m \mathbb{1}_{T_{m-1}<t\leq T_m}\gamma(g(j), m)$$

is the rate $\gamma(g(j), m)$ relevant at time $t$, and $W(t) \sim \mathcal{N}(0, t)$ is standard Brownian motion. Thus, the change in the relative population abundance is modeled using a deterministic term governed by $\gamma_{g(j)}(t)$, which is analogous to the deterministic case, modulated by a stochastic (unpredictable) term governed by $\sigma$.

To model stochastic step-changes, we assume that the step-changes $\Delta_{jm}$ at location $j$ and time $T_m$ follow

$$\Delta_{jm} \sim \mathcal{N}\left(\Delta(g(j), m), \zeta(g(j), m)\sigma^2\right)$$

where the parameters $\zeta(g, m) \in \boldsymbol{\zeta}$ are scaling the stochastic volatility. We assume here that each step change $\Delta_d$ has its unique scaling parameter $\zeta_d$ and hence that $\boldsymbol{\zeta} = \{\zeta_1, \ldots, \zeta_D\}$ and that $\zeta(g, m) = \zeta(g', m')$ whenever $\Delta(g, m) = \Delta(g', m')$.

Under this formulation, the expectation of the individual population trends $\tilde{\varphi}_j(t, \boldsymbol{\theta})$ follow the trend of the deterministic model above:

$$\mathbb{E}[\tilde{\varphi}_j(t_k, \boldsymbol{\theta})] = \varphi_j(t_k, \boldsymbol{\theta}),$$

to which the stochastic model also converges as $\sigma \to 0$.

The probability density $\mathbb{P}\left(\tilde{\varphi}_j(t_k, \boldsymbol{\theta})|\tilde{\varphi}_j(t_{k-1}, \boldsymbol{\theta})\right)$ has the closed form (Karlin and Taylor, 1981)

$$\log\tilde{\varphi}_j(t_k, \boldsymbol{\theta}) = \log\tilde{\varphi}_j(t_{k-1}, \boldsymbol{\theta}) + \epsilon_{jk}; \quad \epsilon_{jk} \sim \mathcal{N}\left(e_{jk}, v_{jk}\right), \tag{3}$$

with the Gaussian mean and variance given by

$$e_{jk} = \sum_{m=1}^{M} \left[\tilde{\rho}_m(t_k)\gamma(g(j), m) + \mathbb{1}_{t_{k-1}<T_m\leq t_k}\Delta(g(j), m)\right] - \frac{\sigma^2}{2}(t_k - t_{k-1}),$$
$$v_{jk} = \sigma^2 \left[(t_k - t_{k-1}) + \sum_{m=1}^{M} \mathbb{1}_{t_{k-1}<T_m\leq t_k}\zeta(g(j), m)\right].$$

Here, we made use of the assumption that observations are affected by step changes occurring at the same time (and hence that any step change at $T_0$ has no detectable effect, see above) and introduced the factors $\tilde{\rho}_m(t_k)$ analogous to eq. 2:

$$\tilde{\rho}_m(t_k) := \begin{cases} T_m - T_{m-1} & \text{if} \quad t_k \geq T_m \text{ and } t_{k-1} \leq T_{m-1}, \\ t_k - T_{m-1} & \text{if} \quad T_{m-1} < t_k < T_m \text{ and } t_{k-1} \leq T_{m-1}, \\ T_m - t_{k-1} & \text{if} \quad t_k \geq T_m \text{ and } T_{m-1} < t_{k-1} < T_m, \\ t_k - t_{k-1} & \text{if} \quad T_{m-1} < t_k < T_m \text{ and } T_{m-1} < t_{k-1} < T_m, \\ 0 & \text{otherwise.} \end{cases}$$

These factors indicate whether the interval between the sampling times $t_{k-1}$ and $t_k$ covers the epoch $m$ fully, partially in the beginning, partially in the end, partially within, or not at all. Accordingly, the rate $\gamma(g(j), m)$ was active for the full length of the epoch $m$, from the start of the epoch until $t_k$, from $t_{k-1}$ until the end of the epoch, from $t_{k-1}$ until $t_k$, or not at all, respectively. In other words:

$$\tilde{\rho}_m(t_k) := \text{length of} \quad [T_{m-1}, T_m] \cap [t_{k-1}, t_k].$$

Note that this model could easily be extended to a correlated Brownian process to account for correlations between locations.

### Modeling observations

Let $\boldsymbol{n}_{ij} = (n_{ij1}, \ldots, n_{ijK})$ denote the observed counts obtained with survey method $i$ at location $j$ during survey times $k = 1, \ldots, K$ at known times $t_k$. We assume these counts are proportional to known efforts $s_{ijk}$ and unknown rates $\lambda_{ij}(t_k)$. The efforts $s_{ijk}$ may represent e.g. the length of a particular transect or camera trap survey, but can also be modeled using covariates (see below). The rates $\lambda_{ij}(t_k)$ describe the rates at which observations are recorded per unit of effort and are modeled as a function of both the population abundance at time $t_k$ and the method- and site-specific detection probability $\delta_{ij}$ as

$$\lambda_{ij}(t_k) = N_{jT_0}\varphi_j(t_k, \boldsymbol{\theta})\delta_{ij}, \tag{4}$$

where $N_{jT_0}$ is the number of individuals at time $T_0$. The relative population abundance $\varphi_j(t_k, \boldsymbol{\theta})$ is given by eq. 1 for the deterministic case and eq. 3 for the stochastic case. Since population abundance and detection probability are confounded, they can not be estimated individually. However, as we show below, by conditioning on the total number of observations, the potentially complex process of detection (Hofmeester et al., 2019) does not need to be modeled.

Similar to Link and Sauer (1997), we model observed counts as following either a Poisson or a negative binomial distribution.

#### Poisson case

Here, we assume that the counts $n_{ijk}$ are Poisson distributed as

$$n_{ijk} \sim \text{Poisson}\left(\lambda_{ij}(t_k)s_{ijk}\right),$$

with rates $\lambda_{ij}(t_k)$ given by eq. 4. To make inferences about the rate parameters $\boldsymbol{\theta}$, we condition on the total number of observations across all survey times $\nu_{ij} = \sum_k n_{ijk}$ (see Link and Sauer, 1997). The conditional distribution of $\boldsymbol{n}_{ij}$ given $\nu_{ij}$ is multinomial (Johnson et al., 1997)

$$\boldsymbol{n}_{ij}|\nu_{ij} \sim \text{Multinom}\left(p_{ij1}, \ldots, p_{ijK}\right)$$

with probabilities

$$p_{ijk} = \frac{\lambda_{ij}(t_k)s_{ijk}}{\sum_{k'}\lambda_{ij}(t_{k'})s_{ijk'}} = \frac{N_{jT_0}\varphi_j(t_k,\boldsymbol{\theta})\delta_{ij}s_{ijk}}{\sum_{k'}N_{jT_0}\varphi_j(t_{k'},\boldsymbol{\theta})\delta_{ij}s_{ijk'}} = \frac{\varphi_j(t_k,\boldsymbol{\theta})s_{ijk}}{\sum_{k'}\varphi_j(t_{k'},\boldsymbol{\theta})s_{ijk'}}. \tag{5}$$

Conditioning on the total number of observations focuses attention on how counts are distributed across survey times rather than on their absolute magnitude. Because both the initial population size ($N_{jT_0}$ ) and the detection probability ($\delta_{ij}$) are assumed constant across time, they act as common scaling factors for all survey occasions. When forming the conditional probabilities, these factors appear in both numerator and denominator and therefore cancel out. As a result, inference about the rate parameters $\gamma$ depends only on the relative changes in counts over time, not on unknown absolute densities or detection probabilities. We relax and discuss our assumption of time-invariant detection probabilities below.

The multinomial likelihood of the full observation vector $\boldsymbol{n} = (\boldsymbol{n}_{11}, \ldots, \boldsymbol{n}_{IJ})$, conditional on $\boldsymbol{\nu} = (\nu_{11}, \ldots, \nu_{IJ})$, is

$$\mathbb{P}(\boldsymbol{n}|\boldsymbol{\theta},\boldsymbol{\nu}) \propto \prod_{i,j,k} p_{ijk}^{n_{ijk}}.$$

We use the non-informative Jeffrey's prior

$$\mathbb{P}(\boldsymbol{\theta}) \propto \sqrt{\det \mathcal{I}(\boldsymbol{\theta})},$$

which is (up to a normalizing constant) the square root of the determinant of the Fisher information matrix

$$\mathcal{I}(\boldsymbol{\theta}) = -\mathbb{E}\left[\frac{\mathrm{d}^2}{\mathrm{d}\boldsymbol{\theta}^2}\log \mathbb{P}(\boldsymbol{n}|\boldsymbol{\theta},\boldsymbol{\nu})\right]$$

Using $\mathbb{E}[n_{ijk}] = \nu_{ij} p_{ijk}$ and $\sum_k p_{ijk} = 1$, we get

$$\begin{aligned}
\mathbb{E}\left[\frac{\partial^2}{\partial\theta_u \partial\theta_v} \log \mathbb{P}(\boldsymbol{n}|\boldsymbol{\theta}, \boldsymbol{\nu})\right] &= \mathbb{E}\left[\sum_{i,j,k} n_{ijk} \frac{\partial^2}{\partial\theta_u \partial\theta_v} \log p_{ijk}\right] \\
&= \sum_{i,j,k} \mathbb{E}[n_{ijk}] \frac{\partial^2}{\partial\theta_u \partial\theta_v} \log p_{ijk} \\
&= \sum_{i,j} \nu_{ij} \sum_k p_{ijk} \frac{\partial^2}{\partial\theta_u \partial\theta_v} \log p_{ijk} \\
&= \sum_{i,j} \nu_{ij} \sum_k p_{ijk} \frac{\partial}{\partial\theta_u} \left(\frac{1}{p_{ijk}} \frac{\partial p_{ijk}}{\partial\theta_v}\right) \\
&= \sum_{i,j} \nu_{ij} \sum_k \left(\frac{\partial^2 p_{ijk}}{\partial\theta_u \partial\theta_v} - \frac{1}{p_{ijk}} \frac{\partial p_{ijk}}{\partial\theta_u} \frac{\partial p_{ijk}}{\partial\theta_v}\right) \\
&= -\sum_{i,j} \nu_{ij} \sum_k \frac{1}{p_{ijk}} \frac{\partial p_{ijk}}{\partial\theta_u} \frac{\partial p_{ijk}}{\partial\theta_v},
\end{aligned}$$

thus we obtain the elements of the Fisher information matrix :

$$\mathcal{I}_{uv}(\boldsymbol{\theta}) = \sum_{i,j} \nu_{ij} \sum_k \frac{1}{p_{ijk}} \frac{\partial p_{ijk}}{\partial\theta_u} \frac{\partial p_{ijk}}{\partial\theta_v}.$$

Another way to write the Fisher matrix is

$$\mathcal{I}(\boldsymbol{\theta}) = \sum_{i,j} \nu_{ij} \mathbf{R}_{ij}' \mathbf{P}_{ij}^{-1} \mathbf{R}_{ij}$$

where $\mathbf{P}_{ij} = \mathrm{diag}(p_{ij1}, \ldots, p_{ijK})$ and $\mathbf{R}_{ij}$ is a $K \times (L + D)$-matrix with elements

$$[\mathbf{R}_{ij}]_{ku} = \frac{\partial p_{ijk}}{\partial\theta_u}.$$

Now remember that the parameter space consists of two types of parameters, the rates $\gamma_\ell$ and the step changes $\Delta_d$ :

$$\boldsymbol{\theta} = (\boldsymbol{\gamma}, \boldsymbol{\Delta}) = (\gamma_1, \ldots, \gamma_L, \Delta_1, \ldots, \Delta_D).$$

Using (5) and (1) we calculate the following partial derivatives:

$$\frac{\partial}{\partial \gamma_l}\varphi_j(t_k,\boldsymbol{\theta}) = \varphi_j(t_k,\boldsymbol{\theta})\sum_m \mathbb{1}_{\gamma(g(j),m)=\gamma_l}\rho_m(t_k),$$

where the sum runs across all epochs for which the rate $\gamma(g(j), m)$ corresponds to the rate $\gamma_l$ and where the terms $\rho_m(t_k)$ are given by (2). Moreover,

$$\frac{\partial p_{ijk}}{\partial \gamma_l} = \frac{\frac{\partial}{\partial \gamma_l}\varphi_j(t_k,\boldsymbol{\theta})s_{ijk}\left(\sum_{k'}\varphi_j(t_{k'},\boldsymbol{\theta})s_{ijk'}\right) - \varphi_j(t_k,\boldsymbol{\theta})s_{ijk}\frac{\partial}{\partial \gamma_l}\left(\sum_{k'}\varphi_j(t_{k'},\boldsymbol{\theta})s_{ijk'}\right)}{\left(\sum_{k'}\varphi_j(t_{k'},\boldsymbol{\theta})s_{ijk'}\right)^2}.$$

Putting these expressions together, we arrive at

$$\frac{\partial p_{ijk}}{\partial \gamma_l} = \frac{\varphi_j(t_k,\boldsymbol{\theta})s_{ijk}}{(\sum_{k'}\varphi_j(t_{k'},\boldsymbol{\theta})s_{ijk'})^2}\sum_{m=1}^{M}\mathbb{1}_{\gamma(g(j),m)=\gamma_l}\sum_{k'}(\rho_m(t_k)-\rho_m(t_{k'}))\varphi_j(t_{k'},\boldsymbol{\theta})s_{ijk'}.$$

In an analogous way, we obtain

$$\frac{\partial p_{ijk}}{\partial \Delta_d} = \frac{\varphi_j(t_k,\boldsymbol{\theta})s_{ijk}}{(\sum_{k'}\varphi_j(t_{k'},\boldsymbol{\theta})s_{ijk'})^2}\sum_{m=1}^{M-1}\mathbb{1}_{\Delta(g(j),m)=\Delta_d}\sum_{k'}(\mathbb{1}_{t_k\geq T_m}-\mathbb{1}_{t'_k\geq T_m})\varphi_j(t_{k'},\boldsymbol{\theta})s_{ijk'}.$$

For the stochastic model, Jeffrey's prior cannot be determined analytically. But since $\mathbb{E}[\tilde{\varphi}_j(t_k,\boldsymbol{\theta})] = \varphi_j(t_k,\boldsymbol{\theta})$, the deterministic solution will serve as a good approximation.

#### Negative Binomial (NB) case

Often, ecological count data violates the Poisson assumption of mean equals variance and this overdispersion is typically modeled using the negative binomial (NB) distribution (e.g. Link and Sauer, 1997; Lindén and Mäntyniemi, 2011). The NB distribution arises naturally when treating the Poisson rates as latent variables that follow a conjugate Gamma distribution. Following Link and Sauer (1997), we choose

$$\tilde{\lambda}_{ij}(t_k) \sim \text{Gamma}\left(\frac{\lambda_{ij}(t_k)s_{ijk}}{a_i}, \frac{1}{a_i}\right),$$
$$n_{ijk} \sim \text{Poisson}\left(\tilde{\lambda}_{ij}(t_k)\right),$$

such that

$$n_{ijk} \sim \mathrm{NB}\left(\frac{\lambda_{ij}(t_k)s_{ijk}}{a_i}, \frac{1}{1+a_i}\right),$$

with a method-specific, constant variance/mean ratio of

$$\frac{\mathrm{Var}(n_{ijk})}{\mathbb{E}[n_{ijk}]} = 1 + a_i,$$

which corresponds to the Poisson case in the limit $a_i \to 0$.

The conditional distribution of $\boldsymbol{n}_{ij}$ given $\nu_{ij}$ is Dirichlet-multinomial (DM, Johnson et al., 1997)

$$\boldsymbol{n}_{ij}|\nu_{ij} \sim \mathrm{DM}\left(\alpha_{ij1}, \ldots, \alpha_{ijK}\right),$$

with parameters

$$\alpha_{ijk} = \frac{\lambda_{ij}(t_k)s_{ijk}}{a_i} = \frac{N_{jT_0}\delta_{ij}}{a_i}\varphi_j(t_k, \boldsymbol{\theta})s_{ijk}. \tag{6}$$

In contrast to the Poisson case, the nuisance parameters $N_{jT_0}$ and $\delta_{ij}$ do not cancel out and have to be inferred along $\boldsymbol{\theta}$ and $a_i$. However, they have little influence on the data as the mean is independent of them:

$$\mathbb{E}[\boldsymbol{n}_{ij}|\nu_{ij}] = \nu_{ij}\boldsymbol{\pi}_{ij} \perp N_{jT_0}, \delta_{ij}, a_i,$$

with probabilities $\boldsymbol{\pi}_{ij} = (\pi_{ij1}, \ldots, \pi_{ijK})'$ identical to the Poisson case (eq. 5):

$$\pi_{ijk} = \frac{\alpha_{ijk}}{\sum_{k'} \alpha_{ijk'}} = \frac{\varphi_j(t_k, \boldsymbol{\theta})s_{ijk}}{\sum_{k'} \varphi_j(t_{k'}, \boldsymbol{\theta})s_{ijk'}} = p_{ijk}.$$

The Dirichlet-multinomial likelihood of the full observation vector $\boldsymbol{n} = (\boldsymbol{n}_{11}, \ldots, \boldsymbol{n}_{IJ})$, conditional on $\boldsymbol{\nu} = (\nu_{11}, \ldots, \nu_{IJ})$, is

$$\mathbb{P}(\boldsymbol{n}|\boldsymbol{\theta}, \boldsymbol{\nu}) \propto \prod_{i,j} \frac{\Gamma(\alpha_{ij0})}{\Gamma(\nu_{ij}\alpha_{ij0})} \prod_{k} \frac{\Gamma(n_{ijk} + \alpha_{ijk})}{\Gamma(\alpha_{ijk})},$$

where $\alpha_{ij0} = \sum_k \alpha_{ijk}$.

Note that $N_{jT_0}, \delta_{ij}$ and $a_i$ are confounded with multiple combinations of values leading to the same Dirichlet-multinomial likelihood. To avoid the resulting non-identifiability issues, we define

$$\frac{N_{jT_0}\delta_{ij}}{a_i} = \frac{\mu_{ij}}{b_i},$$

and infer the vectors $\boldsymbol{\mu} = (\mu_{11}, \ldots, \mu_{IJ})$ and $\boldsymbol{b} = (b_1, \ldots, b_I)$ with the additional constraint $\sum_j \mu_{ij} = 1$.

The prior is of the form $\mathbb{P}(\boldsymbol{\mu}, \boldsymbol{b}, \boldsymbol{\theta}) = \mathbb{P}(\boldsymbol{\mu})\mathbb{P}(\boldsymbol{b}|\boldsymbol{\mu})\mathbb{P}(\boldsymbol{\theta}|\boldsymbol{\mu}, \boldsymbol{b})$, for which Jeffrey's prior can not be determined analytically. Similar to Link and Sauer (1997), however, we can use a quasi-likelihood approximation (Wedderburn, 1974), for which Jeffrey's prior $\mathbb{P}(\boldsymbol{\mu}, \boldsymbol{b}, \boldsymbol{\theta}) \approx \mathbb{P}(\boldsymbol{\mu})\mathbb{P}(\boldsymbol{b})\mathbb{P}(\boldsymbol{\theta})$ with $\mathbb{P}(\boldsymbol{\theta})$ identical to the Poisson case. We further choose the improper prior $\mathbb{P}(\boldsymbol{\mu}) \propto 1$ for all $\boldsymbol{\mu}$ that adhere to the constraint $\sum_j \mu_{ij} = 1$ and $\mathbb{P}(\boldsymbol{\mu}) = 0$ otherwise, and the exponential prior $b_i \sim \text{Exp}(\lambda_b)$, $\mathbb{P}(\boldsymbol{b}) = \prod_i \mathbb{P}(b_i)$ with default rate $\lambda_b = 0.1$.

#### Modeling survey efforts using covariates

Accounting for covariates is straightforward under the models proposed above. Following others (e.g. Link and Sauer, 1997, 1998; Kéry et al., 2009; Fewster et al., 2000), we model the effective efforts as a function of $C_i$ user-provided and method-specific covariates such as, for instance, the number of kilometers walked on a transect. Let $x^{(s)}_{ijkc}$ denote covariate $c = 1, \ldots, C_i$ for method $i$, location $j$ and survey time $k$, and let $\beta^{(s)}_{ic}$ denote the corresponding coefficient. The effort is then modeled as

$$s_{ijk} = \sum_c \beta^{(s)}_{ic} \tilde{x}^{(s)}_{ijkc},$$

where, to avoid non-identifiability issues, we standardize the covariates per method such that

$$\tilde{x}^{(s)}_{ijkc} = JK \frac{x^{(s)}_{ijkc}}{\sum_{j',k'} x^{(s)}_{ij'k'c}},$$

and impose the constraint that $\sum_c \beta_{ic}^{(s)} = 1$. In case the user supplies a single covariate ($C_i = 1$), we set $s_{ijk} = x_{ijk1}^{(s)}$ and do not infer the regression coefficient $\beta_{i1}^{(s)}$.

Note that the covariates $x_{ijkc}^{(s)}$ must be positive and that we do not include an intercept since an actual effort of zero (e.g. walking zero kilometers) will always result in zero counts.

### Relaxing the assumption of time-invariant detection rates

Count data does not allow to distinguish between changes in population abundance and changes in detection rates. In the models introduced above, we assumed that detection rates may vary between location and methods, but are constant through time. As we argue in more detail in the discussion, this assumption is likely appropriate for many ecological surveys. However, it can be relaxed through user-provided covariates.

Let $\delta_{ijk}$ denote the detection probability of method $i$ at location $j$ and survey time $k$. We may model it through $D_i$ method-specific covariates such as, for instance, the experience of the observer, the time of the day or habitat structure. Let $x_{ijkd}^{(\delta)}$ denote covariate $d = 1, \ldots, D_i$, for method $i$, location $j$ and survey time $k$. The detection rate is then modeled as

$$\text{logit } \delta_{ijk} = \beta_{i0}^{(\delta)} + \sum_d \beta_{id}^{(\delta)} \tilde{x}_{ijkd}^{(\delta)},$$

where $\beta_{i0}$ and $\beta_{id}^{(\delta)}$ are method-specific intercepts and coefficients. We standardized the covariates per method such that

$$\tilde{x}_{ijkd}^{(\delta)} = \frac{x_{ijkd}^{(\delta)} - \bar{x}_{id}^{(\delta)}}{\varsigma_{id}},$$

where $\bar{x}_{id}^{(\delta)} = \frac{1}{JK} \sum_{j,k} x_{ijkd}^{(\delta)}$ is the mean across all locations and survey times and $\varsigma_{id}$ their standard deviation. Note that some covariates $x_{ijkd}^{(\delta)}$ may be negative (e.g. temperature).

If time-dependent detection probabilities are modeled through covariates, the relevant probabilities for the Poisson case become (adding $\delta_{ijk}$ to eq. 5 in the main text)

$$p_{ijk} = \frac{\varphi_j(t_k, \boldsymbol{\gamma}) s_{ijk} \delta_{ijk}}{\sum_{k'} \varphi_j(t_{k'}, \boldsymbol{\gamma}) s_{ijk'} \delta_{ijk'}}.$$

For the NB case, the relevant parameters become (replacing $\delta_{ij}$ with $\delta_{ijk}$ in eq. 6 in the main text)

$$\alpha_{ijk} = \frac{N_{jT_0} \delta_{ijk} \varphi_j(t_k, \boldsymbol{\gamma}) s_{ijk}}{a_i},$$

and we now define

$$\frac{N_{jT_0}}{a_i} = \frac{\mu_{ij}}{b_i}$$

with the same constraint $\sum_j \mu_{ij} = 1$ to avoid non-identifiability issues with $b_i$.

### Bayesian inference

Given observed counts $\boldsymbol{n}$, we use Markov chain Monte Carlo (MCMC) methods with Metropolis-Hastings updates and symmetric proposal kernels to obtain posterior samples under the models described above. By default, the MCMC algorithm is run with one chain, 100'000 total iterations, a burn-in of 1'000 iterations, a thinning interval of 10, and numBurnin = 10; these values can all be modified by the user via the same named arguments in the function call. In all models, the inferred parameters include $\boldsymbol{\gamma}$ and $\boldsymbol{\Delta}$. For NB models, we further infer $\boldsymbol{b}$ and $\boldsymbol{\mu}$, and for stochastic models additionally $\sigma$ and the vector $\tilde{\boldsymbol{\varphi}}$ of $\tilde{\varphi}_j(t_k, \boldsymbol{\gamma})$ for all locations and timepoints. In case covariates are used, we also infer the vector of relevant coefficients, here denoted by $\boldsymbol{\beta}$. For fast convergence of the MCMC, we obtain initial estimates of the $\boldsymbol{\gamma}$ parameters using ordinary least squares (see SI). We run the MCMC for $10^4$ iterations as burnin and $10^5$ iterations of sampling.

With the inferred $\boldsymbol{\gamma}$, we can address highly relevant questions for species monitoring and conservation: (i) Is a population increasing or decreasing during epoch $m$? This question is answered by the posterior probabilities $\mathbb{P}(\gamma(g, m) > 0|\boldsymbol{n})$ or $\mathbb{P}(\gamma(g, m) < 0|\boldsymbol{n})$

for the population represented by group $g$. (ii) Did a population trend change at a specific time $T_m$? This question, which arises in BA designs, has so far only been addressed indirectly and under parameter rich models (e.g. Fewster et al., 2000). It can readily be answered by determining the posterior probabilities $\mathbb{P}(\gamma(g,m) > \gamma(g,m+1)|\boldsymbol{n})$ or $\mathbb{P}(\gamma(g,m) < \gamma(g,m+1)|\boldsymbol{n})$. iii) Does the trend differ between two populations during epoch $m$? This question, which arises in CI designs, can be answered by determining the posterior probabilities $\mathbb{P}(\gamma(g,m) > \gamma(g',m)|\boldsymbol{n})$ or $\mathbb{P}(\gamma(g,m) < \gamma(g',m)|\boldsymbol{n})$ for the populations represented by groups $g$ and $g'$, respectively. (iv) Finally, the model is flexible regarding the experimental design and arbitrary rates may be compared. In a BACI design, for instance, the control populations represented by group $g$ share the same rate in both epochs ($\gamma(g,m) = \gamma(g,m+1) = \gamma_1$), while the populations with an intervention represented by group $g'$ share that rate in the first ($\gamma(g',m) = \gamma_1$) but have a potentially different rate in the second epoch ($\gamma(g',m+1) = \gamma_2$). In such a case, the posterior probabilities of interest are $\mathbb{P}(\gamma_1 > \gamma_2|\boldsymbol{n})$ or $\mathbb{P}(\gamma_1 < \gamma_2|\boldsymbol{n})$.

### Initialization of $\boldsymbol{\gamma}$

Prior to running an MCMC inference, we use linear regressions to initialize each $\gamma_l \in \boldsymbol{\gamma}$ as follows. We first identify all consecutive ranges of data points that are informative for a specific $\gamma_l$. These ranges may span multiple consecutive epochs for the data points of a group if the same $\gamma_l$ is relevant for these epochs and that group such as in canonical before-after-control-intervention (BACI) designs. We then build a regression model in which all relevant data ranges share $\gamma_l$ but have their own intercept $\alpha_p, p = 1, \ldots, P$, where $P$ denotes the total of identified ranges,

$$\boldsymbol{y} = \mathbf{X}\boldsymbol{\beta} + \epsilon, \tag{7}$$

with coefficients (intercepts and slope $\gamma_l$)

$$\boldsymbol{\beta} = \begin{pmatrix} \alpha_1 \\ \vdots \\ \alpha_P \\ \gamma_l \end{pmatrix}$$

and

$$\mathbf{X} = \begin{pmatrix} 1 & 0 & 0 & \cdots & 0 & t_{1,1} \\ \vdots & \vdots & \vdots & \ddots & \vdots & \vdots \\ 1 & 0 & 0 & \dots & 0 & t_{1,n_1} \\ 0 & 1 & 0 & \cdots & 0 & t_{2,1} \\ \vdots & \vdots & \vdots & \ddots & \vdots & \vdots \\ 0 & 1 & 0 & \cdots & 0 & t_{2,n_2} \\ \vdots & \vdots & \vdots & \ddots & \vdots & \vdots \\ 0 & 0 & 0 & \cdots & 1 & t_{P,n_P} \end{pmatrix},$$

of which each row corresponds to a data point relevant for $\gamma_l$ and the first $P$ columns indicate the relevant intercept and the last column their survey time. We denote by $t_{p,z}$ the survey time of data point $z$ of range $p$ and by $n_p$ the number of data points in that range. As response variable $\boldsymbol{y}$, we use the log-transformed observed counts per effort for each data point with entries

$$y_{p,z} = \log \frac{n_{p,z}}{s_{p,z}}, \tag{8}$$

where $n_{p,z}$ and $s_{p,z}$ denote the observed counts and sampling efforts of data point $z$ of range $p$, respectively.

The ordinary least squares (OLS) estimator $\hat{\boldsymbol{\beta}}$ of equation 7 is given by

$$\hat{\boldsymbol{\beta}} = (\mathbf{X}^T\mathbf{X})^{-1}\mathbf{X}^T\boldsymbol{y}.$$

Knowing the $\gamma_l$ of an epoch provides information about the rate parameters of neighboring epochs. We thus initialize the $\gamma_l$ iteratively using the following scheme:

1. We start with the not yet initialized $\gamma_l$ for which we have the most information, which we identify as the $\gamma_l$ with the highest score $\mathcal{S}_l$ calculated as the sum of scores across all ranges relevant for $\gamma_l$, i.e. $\mathcal{S}_l = \sum_p \mathcal{S}_{lp}$. Each range is given a score as follows:

   - A range with less than two data points is given a score $\mathcal{S}_{lp} = 0$.
   - A range with at least two data points is given a score

   $$\mathcal{S}_{lp} = \sum_z \left[ \frac{1}{2} \mathcal{I}(n_{p,c} = 0) + \mathcal{I}(n_{p,c} > 0) \right],$$

     where the sum runs across all data points in range $p$ and $\mathcal{I}$ is an indicator function. Hence, a non-zero data point is worth a score of one and a zero data-point a score of one half.

2. If the highest score is zero, then the system is not identifiable and an error is thrown (e.g. if for no range more than one data point is available).

3. We estimate the coefficients $\hat{\boldsymbol{\beta}} = (\hat{\boldsymbol{\alpha}}, \hat{\gamma}_l)$ using OLS as described above and use the value $\hat{\gamma}_l$ to initialize $\gamma_l$. Since the logarithm of $n_{p,z} = 0$ in equation 8 results in minus infinity, we exclude data points with zero observed counts from the regression. In case excluding such data points implies that $\gamma_l$ cannot be inferred, we set it to zero.

4. We then use the gained knowledge to augment the information of neighboring ranges by predicting the response value at the start and end of each range $p$ relevant for $\gamma_l$ as $\boldsymbol{y}_{T_p}(\hat{\boldsymbol{\alpha}}, \hat{\gamma}_l) = \hat{\boldsymbol{\alpha}} + \mathbb{1}(\hat{\gamma}_l T_p)$, where $\mathbb{1}$ is a vector of ones of length $P+1$ and $T_p$ denotes the time of start or end of range $p$. The predicted response is stored as "data point" and used to initialize other $\gamma_l$, unless it coincides with an existing time point.

5. We then repeat these steps until all $\gamma_l$ have been initialized.

#### Distinguishing between Poisson and negative binomial models

The use of MCMC methods does not lend itself easily to determine whether a Poisson or NB model should be used via Bayesian model selection. To overcome this limitation, we implemented the following simulation-based approach to check whether the inferred overdispersion is outside the range of values expected under a Poisson model: For a given data set $\boldsymbol{n}$, we first infer all parameters under the NB model. Let $\theta_{nb}$ denote that set of inferred parameters, of which we denote the inferred overdispersion parameter $\boldsymbol{b}$ as $\boldsymbol{b}^*$. We then perform $R$ simulations under the conditional Poisson model (i.e. the multinomial distribution conditioned on the observed counts $\nu_{ij}$) using the subset of inferred parameters $\theta_p \subset \theta_{nb}$ relevant for the Poisson model. This includes $\boldsymbol{\gamma}$, but also $\boldsymbol{\beta}$ in case of covariates as well as $\sigma$ and $\tilde{\boldsymbol{\varphi}}$ in case of a stochastic model. For each simulation $r = 1, \ldots, R$, we infer parameters under the NB model and store the posterior mean estimates of $\boldsymbol{b}$ as $\boldsymbol{b}_r$. We then compare the value $\boldsymbol{b}^*$ obtained with the real data against the values $\boldsymbol{b}_1, \ldots, \boldsymbol{b}_R$ obtained under Poisson simulated data and compute the method-specific p-value $p_i = \frac{1}{R}\sum_r \mathbb{1}_{b_{ri} > b_i^*}$ as the fraction of Poisson simulations for which the overdispersion parameter was inferred to be larger than that inferred on the real data. We recommend to use the NB model whenever $p_i$ is below a chosen threshold (e.g. 0.05) and hence the Poisson model is rejected for at least one method $i$.

#### Performance against simulations

To assess the performance of `birp`, we generated simulations under various conditions detailed below. We conducted 1,000 replicate simulations in all cases and repeated the few simulations that resulted in zero counts for all time points. In case of single group, single epoch trends, we only simulated negative or constant trends and determined the power of `birp` to detect the directionality of a trend through the fraction of all replicates for which the posterior $\mathbb{P}(\gamma < 0|\boldsymbol{n}) > 0.95$ if we simulated $\gamma < 0$ and the fraction for which $\mathbb{P}(\gamma < 0|\boldsymbol{n}) > 0.975$ or $\mathbb{P}(\gamma > 0|\boldsymbol{n}) > 0.975$ if we simulated $\gamma = 0$. In case of BA, CI and BACI designs, we used the fraction of replicates for which $\mathbb{P}(\gamma_1 < \gamma_2|\boldsymbol{n}) > 0.95$ if we simulated $\gamma_1 < \gamma_2$, the fraction for which $\mathbb{P}(\gamma_1 > \gamma_2|\boldsymbol{n}) > 0.95$ if we simulated $\gamma_1 > \gamma_2$ and the fraction for which $\mathbb{P}(\gamma_1 < \gamma_2|\boldsymbol{n}) > 0.975$ or $\mathbb{P}(\gamma_1 > \gamma_2|\boldsymbol{n}) > 0.975$ if we simulated $\gamma_1 = \gamma_2$. To model the number of available observations, we varied the expected number of observations at the first time point $\mathbb{E}[n_{ij1}] = N_{jT_0}\delta_{ij}s_{ij1}$. Since only the product of $N_{jT_0}$, $\delta_{ij}$ and $s_{ij1}$ is relevant, we set the detection probabilities $\delta_{ij} = 1$ in all cases, drew the efforts $s_{ijk} \sim \Gamma(1, 2)$ from a Gamma distribution and scaled these such that the average effort was 1.0, and set $N_{jT_0}$ to either 10 or 50.

#### Power to detect trends in a single epoch

First, we examined the effect of the number of time points, the number of available observations, and the number of locations on the power to detect a trend in a single group in a single epoch with $\gamma(1, 1) = \gamma$ under the deterministic Poisson model. As expected, the power of `birp` was higher if the trend was stronger (i.e. $\gamma$ was more negative), if more time points were surveyed, or if more observations were obtained per survey (Figure S1A). However, power was generally high even for limited data. For $J = 5$ locations and $K = 2$ survey times, for instance, the deterministic Poisson model correctly estimated the simulated trends in $> 95\%$ of the cases if $\gamma \leq -0.5$ and if an expected number of $N_{jT_0} = 50$ observations were simulated for the first time point (Figure S1A). When using $K = 5$ survey times, the same $> 95\%$ power was reached with $\gamma \leq -0.2$, already with $N_{jT_0} = 10$. These simulations further show that the posterior probabilities reported by `birp` are well calibrated: if no trend was simulated (i.e. $\gamma = 0$), no trend was detected in 94.7% of all simulations across all cases, close to the 95% expected under the chosen posterior threshold. We next investigated the effect of surveying additional locations in the above setting while fixing $\gamma = -0.2$. As expected, the power to detect a negative trend increased sharply with the number of locations surveyed (Figure S1B). With $J = 10$ locations or more, maximum power was reached if surveys were conducted with $K = 5$

and $N_{jT_0} = 10$.

Second, we explored the effect of increased variation under NB models (varying $a$) and the stochastic model (varying $\sigma$) with $J = 5$, $K = 5$, $N_{jT_0} = 10$ and $\gamma(1,1) = \gamma$. We further used these simulations to explore the effect of incorrectly assuming a Poisson distribution or a deterministic model. While NB/stochastic models account for overdispersion/randomness, more parameters need to be estimated and the data is generally less informative than under the Poisson/deterministic case. Consequently, for the NB case, the power to detect trends decreased with increasing dispersion parameter $a$ (Figure S1C). To investigate the impact of the Poisson assumption, we also re-estimated the parameters under the Poisson model (blue lines in Figure S1C), which led to increased power, at the cost of considerable false-positives at $\gamma = 0$ in case of large overdispersion ($a \geq 1$). For the stochastic case, power decreased with increasing volatility $\sigma$ (Figure S1D). For a small volatility of $\sigma = 0.1$ and using $\gamma = -0.2$, for instance, `birp` correctly estimated the simulated trends in 79% of the cases. For a larger volatility of $\sigma = 0.5$ and $\sigma = 1$, the power dropped below 74% and 40%, respectively, even for $\gamma = -1$. To check for the importance of accounting for variation in local trends, we inferred trends under the deterministic model on the data simulated under the stochastic model. As shown in Figure S1D, a violation of the assumption of a deterministic trend leads to generally higher power, at the cots of many false-positives at $\gamma = 0$, particularly if volatility was high (e.g. 78% in case of $\sigma = 1$). If volatility was small (e.g. $\sigma = 0.1$), however, the benefit of inferring fewer parameters seemed to outweigh the violation of the deterministic assumption as we observed a gain in power without an elevation of false-positives.

#### Testing for the Poisson assumption

The above simulations show that while using Poisson models on overdispersed data leads to increased false positives, they have higher power in case the Poisson assumption is justified. To guide the choice of the appropriate model, we propose a simulation-based approach that evaluates whether the inferred overdispersion parameter differs significantly from what would be expected under a Poisson model (Figure S5A). To assess the power of this approach, we generated simulations under both deterministic Poisson and NB models with varying overdispersion for a single epoch and a single group, using $J = 5$ locations, $K = 5$ time points and $N_{jT_0} = 10$. For each simulation, we then performed inference under i) the Poisson model, ii) the NB model, and iii) the model identified as appropriate based on the simulation-based test. These simulations confirmed the elevated false-positive rates when inferring under a Poisson model from overdispersed data simulated with no trend ($\gamma = 0$, Figure S5B). However, the inference was well calibrated when inferring under a NB model, albeit at the cost of lower statistical power (Figure S5C). When deciding on the appropriate model individually for each simulation (Figure S5D), power could be increased (particularly when overdispersion was small) without any excess in false-positives. Of all simulations conducted with $a = 0.1$, for instance, the Poisson model was deemed appropriate in 93% cases, which resulted in an increase in power to detect a negative trend from 37.6% to 60.5% for $\gamma = -0.1$. For simulations conducted with $a = 1$, $a = 10$ or $a = 100$, in contrast, the Poisson model was deemed appropriate for only 50%, 2% and 1% cases, respectively.

#### Power to detect changes in trends

We next investigated the ability of birp to detect changes in population trends by simulating data under canonical before-after (BA), control-intervention (CI) and before-after-control-intervention (BACI) designs. For each design, we varied the number of time points and rates for both epochs as well as the number of available observations. We focused on deterministic Poisson models in all cases.

- For the BA design, we used two epochs, one group and $\gamma(1, 1) = \gamma_1$ and $\gamma(1, 2) = \gamma_2$. Power was generally higher if more time points were available, if more observations were available, and if the rates $\gamma(1, 1) = \gamma_1$ and $\gamma(1, 2) = \gamma_2$ of the two epochs differed more substantially (Figure S2, first row). In case of five time points in each epoch (5/5 case) and $\gamma_1 = 0$ and $\gamma_2 = -1$, for instance, a decrease in trend was detected with $\mathbb{P}(\gamma_2 < \gamma_1|\boldsymbol{n}) > 0.95$ in 94% of the simulated replicates. If $\gamma_1 = -0.2$, however, that power dropped to 49%. The simulations also show the importance of having multiple time points per epoch. The power was, for instance, substantially higher if two time points were surveyed in each epoch (2/2 case), rather than two before and only one after the time of change $T_1$ (2/1 case), and notably also higher than if five points before and one after were surveyed (5/1 case), despite the higher total number of time points. That said, our simulations show that strong changes in trends can be detected with rather few time points: In the 2/1 case, the power to detect a change from $\gamma_1 = 0$ to $\gamma_2 = -1$ was still 27% for $N_{jT_0} = 50$. Since these simulations were conducted for a single location, we expect much higher power for real world scenarios, as shown for the single epoch case (Figure S1B) above. Finally, the simulations again show that `birp` posterior probabilities are well calibrated: Across all cases in which we simulated no change in trend, a change was detected in exactly the expected 5% of all simulations.

- For the CI design, we focused on cases with two or five time points in a single epoch, surveyed for one control and one intervention location with rates $\gamma(1,1) = \gamma_1$ and $\gamma(2,1) = \gamma_2$ (Figure S2, second row). In case of $\gamma_1 = 0$, power was very comparable but slightly (about 5%) lower than under the BA cases with equal number of surveyed time points. For $N_{jT_0} = 10$ and $\gamma_2 = -1$, for instance, `birp` detected a change in 94% of simulated BA cases with five time points per epoch (5/5 case), but only in 90% of simulated CI cases with five time points per group (-/5 case). From a statistical point of view, these BA and CI designs are indeed very similar, since in both cases two rates have to be inferred, but the BA case can take advantage of an additional comparison, namely the one across the time of change. In case of $\gamma_1 = -0.2$, however, the power was generally higher under comparable CI than BA designs in our simulations. This is because in the CI design as simulated here, both control and intervention locations started with the same expected number of observations at the first time point, while in the BA design the expected number of observations is already reduced at the beginning of the second epoch.

- For the BACI design, we used two locations, under which the control location had the same rate in both epochs ($\gamma(1,1) = \gamma(1,2) = \gamma_1$), while the intervention location had a potentially different rate in the second epoch ($\gamma(2,1) = \gamma_1$, $\gamma(2,2) = \gamma_2$, Figure S2, third row). Under BACI designs, the power to detect changes in trends was higher than under either BA or CI designs due to the additional available data. For the 2/2 case with $\gamma_1 = 0$, $\gamma_2 = -1$ and $N_{jT_0} = 10$, for instance, the power doubled from 23% under a BA design to 46% under a BACI design. For the same parameters but for the 2/1 case, the power increase was from 12% to 18%. Interestingly, the picture was less clear when both design had the same number of surveys. The 2/2 BA case, for instance, had a higher power (23%) than the 2/1 BACI case (18%). This pattern was observed for all but one combinations of $\gamma_1$ and $\gamma_2$ tested and suggests that under the conditions explored here, additional information for $\gamma_2$, as provided under BA designs, may be more relevant than additional information for $\gamma_1$, as provided under BACI designs.

#### All-zeroes and time lags

Finally, we explored the power of `birp` to detect changes under BA designs in the case of local extinctions and in case an intervention affects a population with a time lag. Wauchope et al. (2021) recently identified two cases under which the detection of changes in trends are complicated. The first case, termed *all-zeroes*, describes a situation under which the species went locally extinct and hence only zero counts were obtained for the second epoch. Such cases are detrimental to approaches that compare trends inferred independently for each epoch, as all zero counts do not allow for trend estimation or are interpreted as a stable trend. We here explored the power to detect such changes under the `birp` model, which also exploits the comparison across epochs. As shown in Figure S3AB, `birp` has no issues with all zeroes cases and retains high power. Importantly, however, the power strongly depends on the trend during the first epoch, since strongly negative trends lead to populations that are already close to extinction before the time of change. Since all our simulations start with $N_{jT_0} = 10$, that is particularly visible in the the 5/1 and 5/5 cases, for which the expected number of counts at the time of change $T_1$ is 0.11 in case of $\gamma_1 = -1$, and hence already very close to 0.0.

The second case with complications raised by Wauchope et al. (2021) was a situation under which the impact of an intervention manifested after some time lag only. While `birp` could be readily extended to also infer the most likely time of change, we here explored the power under time lag scenarios under the 5/5 case with $\gamma_1 = 0$. As shown in Figure S3CD, and as expected, long time lags diminish the power to detect changes in trends much more than short time lags. For the case of $\gamma_2 = -1$, for instance, the power dropped from 94% in case of no time lag to 91%, 76% and 15% in case the effect lagged by one, two or four survey seasons, respectively. Importantly, and across all values of $\gamma_2$ tested, the model appears robust to misspecifications of one survey season.

#### Analyses regarding case study 1: Temporal downsampling

Tasmanian devils were previously reported to have declined across almost their entire geographical range in the last three decades after the emergence of the highly contagious devil facial tumour disease (DFTD, Lazenby et al., 2018; Cunningham et al., 2021; Volery et al., 2023). Using `birp` on transect count data, we tested whether the emergence of DFTD resulted in more negative population trends in each Tasmanian bioregion.

As shown in Figure ??A, annual counts varied greatly over time and between bioregions. Consequently, and while our model detected a change to a more negative population trend for five out of six bioregions, results varied in the certainty depending on how many surveys were used (Figure ??C). We found the strongest support for a change to a more negative trend for the bioregion Ben Lomond, for which the posterior probability $\mathbb{P}(\gamma_1 > \gamma_2) > 0.99$ whenever we used three or more surveys after the time of change $T_1$. For the bioregion Northern slopes, we also found support for such a change with

$\mathbb{P}(\gamma_1 > \gamma_2) > 0.95$ if tested using three or four surveys after $T_1$, but with greater uncertainty if more surveys were used. For the bioregions Central Highlands and Southern Ranges, in contrast, we found considerable support for a more negative trend only if five or six surveys after $T_1$ were used, respectively. For the bioregion South East, we identified strong support for a more negative trend with $\mathbb{P}(\gamma_1 > \gamma_2) > 0.99$ right after $T_1$, but not when using additional surveys. For the bioregion King, finally, we estimated a lower population density before $T_1$.

This variation in results is both compatible with a visual inspection of the data as well as previous analyses. For the bioregion King, for instance, reported counts were monotonously increasing for four surveys starting from the one just before $T_1$ and culminating in the third survey after $T_1$, for which the highest counts across the entire time series was reported. There is thus little evidence for a negative effect of DFTD, presumably because the disease did not spread into the entire region (Lazenby et al., 2018). In case of the Southern Ranges, the highest counts of the time series were also reported two years after $T_1$, and the decline consequently only manifested with some delay. In contrast, the counts for South East showed high variability already prior to $T_1$, then a major drop right after $T_1$, followed by an apparent recovery. Consequently, a more negative trend was only inferred when using a single survey after $T_1$. It thus appears as if the devils were differently affected by DFTD in these bioregions, and their trends also likely affected by additional, more local factors.

The region with the clearest signal of an effect was Ben Lomond, for which we inferred strongly different trends before and after $T_1$. Volery et al. (2023) previously reported strong impact of DFTD also for the bioregion Northern Slopes from survey four after $T_1$ until the end of the series. Here, we found strong support when using three or four surveys after $T_1$, but not when using longer time-series. There is, however, a fundamental difference in the quantities that were estimated: `birp` estimates population trends, whereas Volery et al. (2023) estimated impact as the per year population size reduction compared to the expectation in the absence of the disease, a signal that may remain detectable even after a trend stabilized or reversed. We argue that both serve important purposes: while the latter may be used to showcase the impact of a past event, trends are informative about the trajectory of a population and hence the need of further interventions.

### Analyses regarding case study 2: Heterogeneous survey methods and irregular sampling

#### Considered species

We focused on the four apex predators, African wild dog (*Lycaon pictus*), Northern lion (*Panthera leo ssp. leo*), spotted hyena (*Crocuta crocuta*) and African leopard (*Panthera pardus ssp. pardus*), for which data from the Chinko was available. We complemented this data set with data from six mammalian herbivores, namely defassa waterbuck (*Kobus ellipsiprymnus ssp. defassa*), Eastern giant eland (*Tragelaphus derbianus ssp. gigas*), lelwel hartebeest (*Alcelaphus buselaphus ssp. lelwel*), lowland bongo (*Tragelaphus eurycerus ssp. eurycerus*), Western roan antelope (*Hippotragus equinus ssp. koba*) and Western Central African buffalo (*Syncerus caffer ssp. nanus*).

We combined the track count data with data from camera trapping surveys collected from 2012 to 2019 from Aebischer et al. (2020). Detection probabilities most likely varied between locations and seasons (Ait Kaci Azzou et al., 2021). To make observations comparable, we only compared data from camera traps that were installed at the same location (within 40 meters) across different years and considered images taken during the dry (November - April) and wet (May - October) seasons as separate methods. We included consecutive observations of the same species if they were obtained more than 10 minutes apart. The efforts represent the time span during which the camera trap was active, after cutting off 30 minutes at the beginning and at the end of each trapping session to minimise the influence of human presence in the data.

#### Construction of a road network

To make the track counts comparable across years, we used Feature Manipulate Engineering (FME) software to compute a routable road network based on the GPS-tracks recorded during the surveys. We used buffers of 100 meters to aggregate parallel road sections and used topological skeletonization to extract the center line from these aggregated buffers. Once extracted, we created the road network by identifying road intersections. Thus, we divided the Chinko roads into separate segments such that segments never included intersections and were always surveyed in full. We manually verified and corrected the resulting network at nine locations where connections between close segments were missing. We further imposed a maximal segment length of 500 meters and split longer segments accordingly.

To quantify observations, we mapped all observations to their nearest segment in the network, but discarded all observations that were further than 250 meters away from any segment. To quantify efforts, we used the function `shortest_paths` of the package `iGraph` (Csardi and Nepusz, 2006) in `R` (R Core Team, 2024) to identify the shortest path in the network that connected all observations from a set of consecutive days, but considered observations more than 20 kilometers apart to result from independent surveys, thus accounting for independent teams surveying on the same day. We then recorded the number of tracks obtained per segment, with segment-specific efforts reflecting how often a specific segment was surveyed in a given year.

#### Detailed results per species

For most species, our inference indicated a currently positive growth rate and also a positive change in trend following law-enforcement, albeit with variable posterior support (Figure ??). For three out of the six prey species, namely African buffaloes (1,300 observations across all methods and years), waterbucks (961 observations) and giant elands (375 observations), posterior support for a positive change was $\mathbb{P}(\gamma_1 < \gamma_2|\boldsymbol{n}) = 1$, and only slightly lower for hartebeests (0.99, 188 observations) and roan antelopes (0.93, 698 observations). Of those, we inferred that the large ungulates giant eland and hartebeest are currently growing at the fastest rates with posterior mean estimates for $\gamma_2$ at 0.46 and 0.41, respectively. In contrast, waterbucks, African buffaloes and roan antelopes were estimated to grow with rates 0.25, 0.21 and 0.16, respectively. No change in trend was inferred for bongos (233 observations), for which Aebischer et al. (2020) speculated that due to their use of close-canopy forests and their elusive behavior, they were barely affected by direct poaching by herders. However, we inferred a currently growing population with $\mathbb{P}(\gamma_2 > 0|\boldsymbol{n}) = 0.88$ at a rate of 0.23 also for that species.

We also found posterior support with $\mathbb{P}(\gamma_1 < \gamma_2|\boldsymbol{n}) > 0.9$ for a positive change for three out of the four predator species, namely for African leopard (0.96, 288 observations), African wild dog (0.92, 36 observations) and Northern lion (0.91, 38 observations), albeit with a larger uncertainty regarding the exact growth rates for wild dogs and lions due to their small number of observations. All three species were inferred to currently grow with posterior mean estimates for $\gamma_2$ at 0.20, 0.17 and 0.27, respectively. For hyenas, which had the largest number of observations (540), we only found moderate support for both a currently growing population ($\mathbb{P}(\gamma_2 > 0|\boldsymbol{n}) = 0.78$) as well as a positive change in trend since 2017 ($\mathbb{P}(\gamma_1 < \gamma_2|\boldsymbol{n}) = 0.73$). This may reflect that in contrast to lions, hyenas were not directly targeted by poaching herders.

### Supplementary Figures

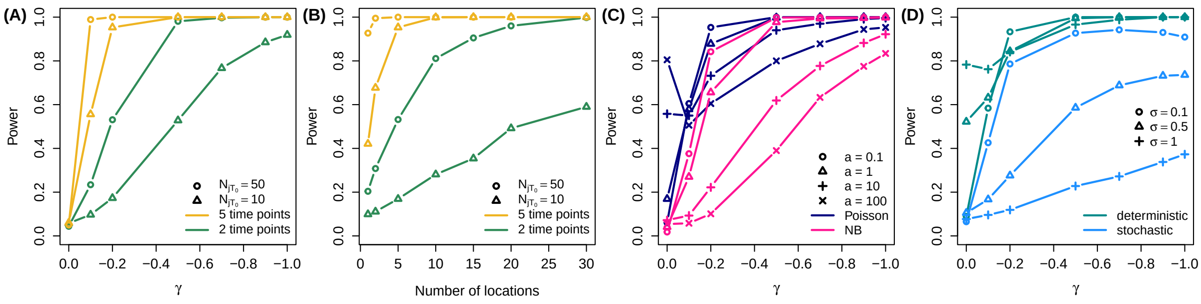


**Fig. S1:** Simulation results for a single epoch and group. Shown are the fraction of 1000 replicates for which the model correctly inferred the directionality of the negative trend with $> 0.95$ posterior support. For the case of $\gamma = 0$, the power was calculated as the fraction of replicates for which any change was inferred with that posterior support. (A) A deterministic Poisson model with $J = 5$ locations for different values of $\gamma$, $K = 2$ or $K = 5$ time points and $N_{jT_0} = 10$ or $N_{jT_0} = 50$ expected observations at the first time point. (B) A deterministic Poisson model as in A but for a fixed $\gamma = -0.2$ and different number of locations. (C) A deterministic negative binomial (NB) model with $J = 5$ locations for different values of $\gamma$ and overdispersion parameter $a$. The pink and blue lines correspond to inference under the NB and Poisson model, respectively, on the same simulations. (D) A stochastic Poisson model with $J = 5$ locations for different values of $\gamma$ and volatility parameter $\sigma$. The light blue and green lines correspond to inference under a stochastic and deterministic Poisson model, respectively, on the same simulations.

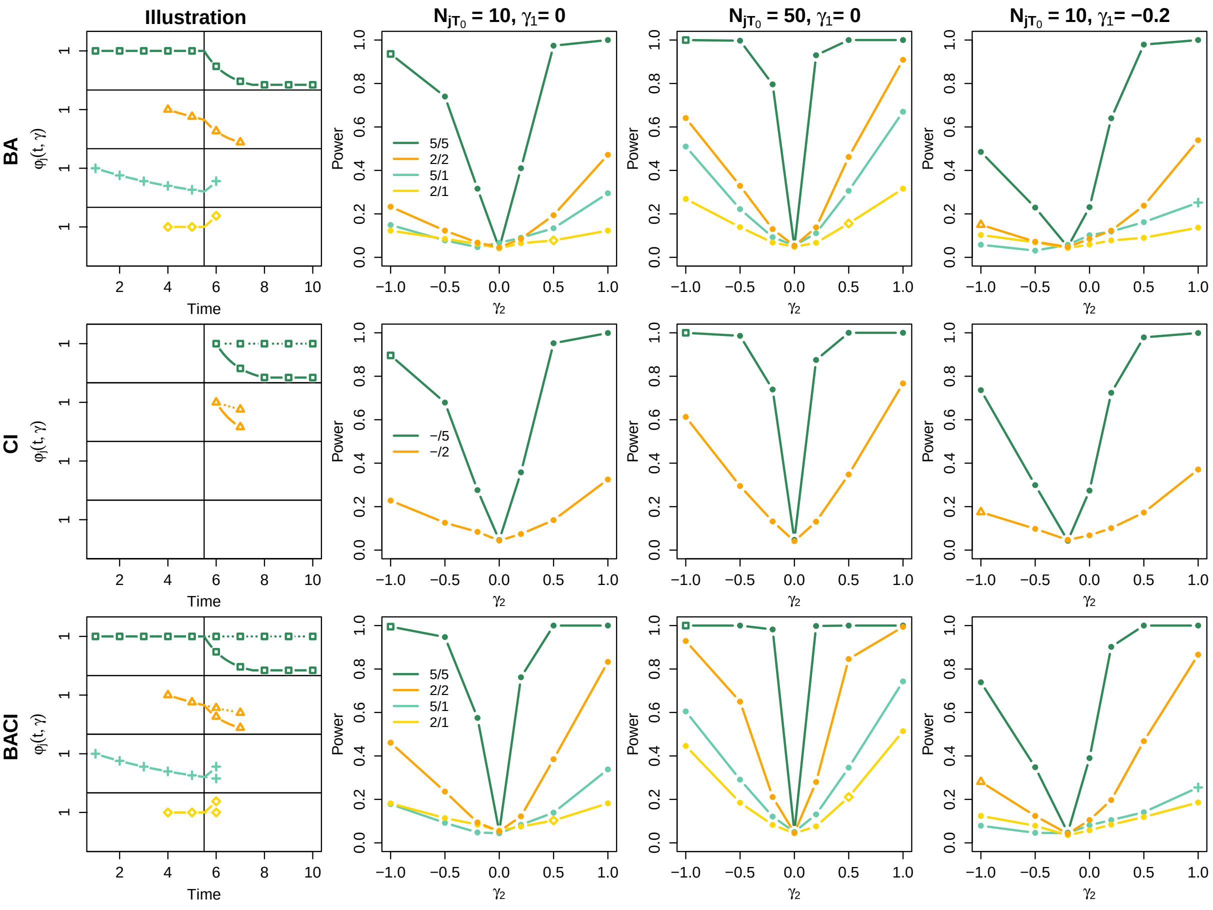


**Fig. S2:** Simulation results for canonical before-after (BA, first row), control-intervention (CI, second row) and before-after-control-intervention designs (BACI, third row) for different values of $\gamma_2$ in the second epoch or intervention group (x-axis) and for different numbers of survey times (colors). Columns differ in the expected number of observations $N_{jT_0}$ at the first time point and the value of $\gamma_1$. For each design, the first column shows an illustration of the population trend $\varphi_j(t, \boldsymbol{\theta})$ for four specific simulation settings, indicated by the matching color and plotting symbol. In case of CI and BACI designs, the dotted lines represent the trend of the control group. The reported power refers to the fraction of 1000 replicates for which the model correctly inferred the directionality of the change with $> 0.95$ posterior support. For the cases of $\gamma_1 = \gamma_2$, we report the fraction of replicates for which any change was inferred with that posterior support.

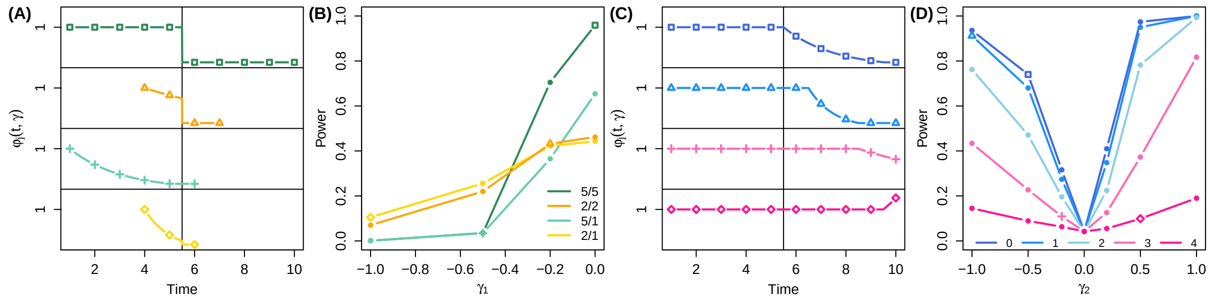


**Fig. S3:** Simulation results for cases of local extinctions (A,B) and time lags (C,D). Panels A and C show the population trend $\varphi_j(t, \boldsymbol{\theta})$ for four specific simulation settings, indicated by the matching color and plotting symbol. Power was calculated as for Figure S2. (A, B) All-zeroes scenario where a constant or negative population trend in the first epoch ($\gamma_1$) is followed by a local extinction event resulting in all-zero counts in the second epoch. We varied the rates of the first epoch $\gamma_1$ (x-axis in B) as well as the number of survey times (colors). (C, D) Time lag scenario where the impact of an intervention manifests after some time only. We varied the rates of the second epoch $\gamma_2$ (x-axis in D) as well as the magnitude of the time lag (colors).

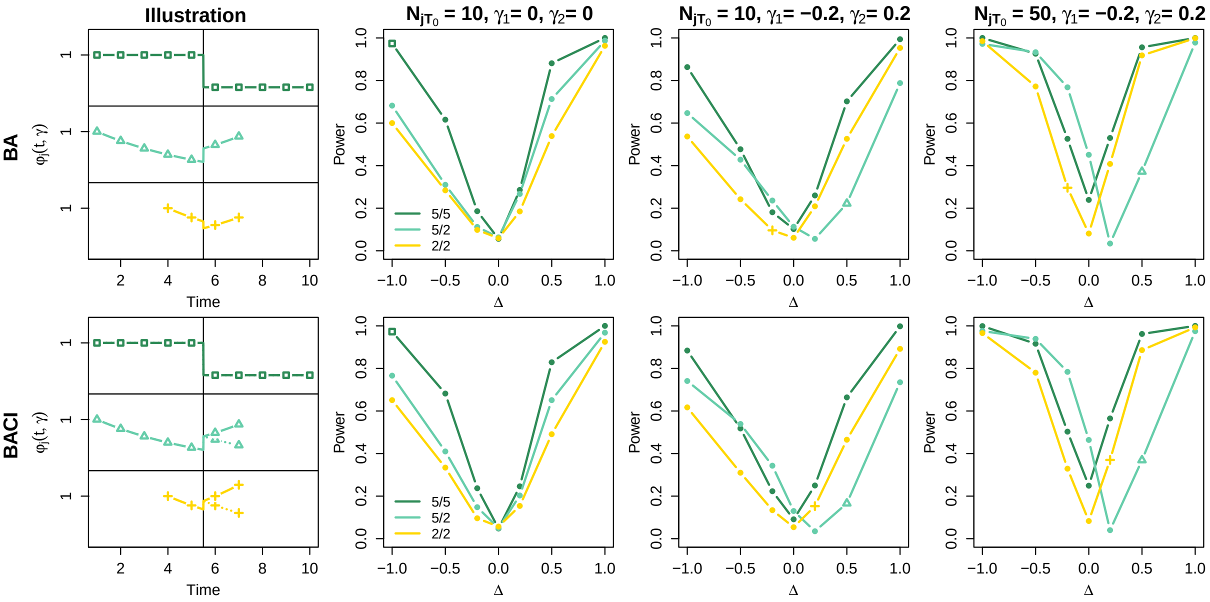


**Fig. S4:** Simulation results for before-after (BA, first row) and before-after-control-intervention designs (BACI, second row) for different values of $\Delta$ (step change) between the first and second epoch (x-axis) and for different numbers of survey times (colors). Columns differ in the expected number of observations $N_{jT_0}$ at the first time point and the value of $\gamma_1$ and $\gamma_2$. For each design, the first column shows an illustration of the population trend $\varphi_j(t, \boldsymbol{\theta})$ for three specific simulation settings, indicated by the matching color and plotting symbol. In the BACI design, the dotted lines represent the trend of the control group. The reported power refers to the fraction of 1000 replicates for which the model correctly inferred the directionality of the change with $> 0.95$ posterior support. For the cases of $\Delta = 0$, we report the fraction of replicates for which any change was inferred with that posterior support.

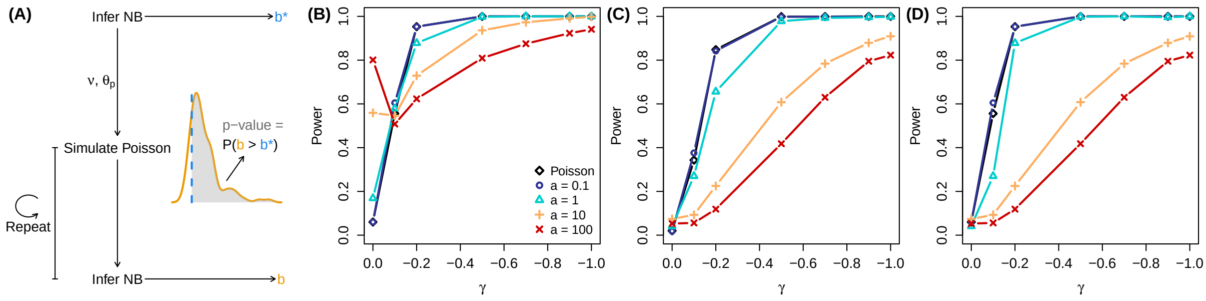


**Fig. S5:** Simulation results for distinguishing between Poisson and negative binomial (NB) models. (A) Scheme illustrating the simulation-based approach used to assess whether the inferred overdispersion $b^*$ (blue) is outside the range of values $b$ expected under a Poisson model (orange). If the p-value is smaller than some threshold (e.g. 0.05), the Poisson model is rejected and the data should be analyzed using a NB model. (B,C,D) Simulations for $J = 5$ locations, different values of $\gamma$ (x-axis) and under deterministic Poisson models (black diamonds) or deterministic NB models with different values of the overdispersion parameter $a$ (colors). Power was calculated as for Figure S1. (B) Inference under a Poisson model. (C) Inference under a NB model. (D) Inference under the model deemed appropriate by the simulation-based approach.
